# System Identification and Control for Optogenetics in Mammalian Nucleocytoplasmic Transport

**DOI:** 10.64898/2026.06.26.734178

**Authors:** Menno van Laarhoven, Alfredo Rates, Josiah B. Passmore, Shengling Shi, Ihor Smal, Lukas C. Kapitein, Carlas S. Smith

## Abstract

Optogenetics enables experiments in out-of-equilibrium conditions to clarify biological mechanisms and quantify biophysical parameters. However, modelling and control techniques to study mammalian cell biology under optogenetic perturbation remain underutilised. Here, we benchmark these methods within mammalian cells by steering nucleocytoplasmic transport via the optogenetic LEXY protein in outcome-driven microscopy. First, we employ system identification to obtain models that predict transport dynamics by minimising the prediction error. We quantify this prediction accuracy for one biophysical model and two black-box models. Second, we evaluate closed-loop control efficacy by steering transport along a predefined trajectory using model-free Proportional Integral (PI) control, model-based Linear Quadratic Regulation (LQR) and Model Predictive Control (MPC). Both the predictive models and the applied control techniques demonstrate robust performance against cell-to-cell variation. This biological variation is quantified by the parameter distributions obtained from model identification with single-cell trajectories. While we show that model-free techniques such as PI and gain-scheduled PI achieve steering without explict model knowledge, predictive architectures offer better performance under this cell-to-cell variation and time-varying setpoints. Moreover, black-box predictive accuracy suggests that this model-based control is possible, even when explicit mechanistic understanding is missing. Ultimately, we demonstrate that predictive modelling and optogenetics enable quantitative characterisation and precise manipulation of mammalian cells, while offering practical guidelines for the implementation of these techniques.

**SIGNIFICANCE:** Optogenetics provides the unique ability to steer cellular processes with light. Variable and localised light inputs allow us to extract more data and guide living cells to specific states. Despite this potential, application of predictive modelling and control techniques with optogenetics remain underutilised in mammalian cell biology. Here, we benchmark these techniques within mammalian cells to steer nucleocytoplasmic transport. Benchmarking these techniques demonstrates their direct applicability to mammalian systems, enabling the extraction of new biological insight in outcome-driven microscopy.

## INTRODUCTION

Fluorescence microscopy serves as a cornerstone of modern cell biology, enabling the visualisation of intracellular dynamics with high resolution. However, traditional microscopes act as passive observation tools that cannot adapt to evolving conditions. Smart microscopy adjusts to these conditions by changing acquisition parameters in real time to, for example, minimise phototoxicity or track objects (1, 2). Yet, while these platforms adapt to the sample, they still treat biological samples as passive entities. Pairing microscopy with optogenetics changes this paradigm by allowing us to control cellular processes with light (3–6). Instead of simple on/off experiment designs, variable and localised light inputs allow us to guide cells to specific states and extract more data (7, 8). More specifically, dynamic inputs enable guidance of cellular processes for experiments in complex out-of-equilibrium conditions to quantify biophysical parameters and clarify biological mechanisms. To achieve these dosed and localised perturbations, optogenetics is often paired with a smart microscopy pipeline and a light modulation device, enabling outcome-driven microscopy (Figs. 1a and 1f) (9–13).

**Figure 1.**
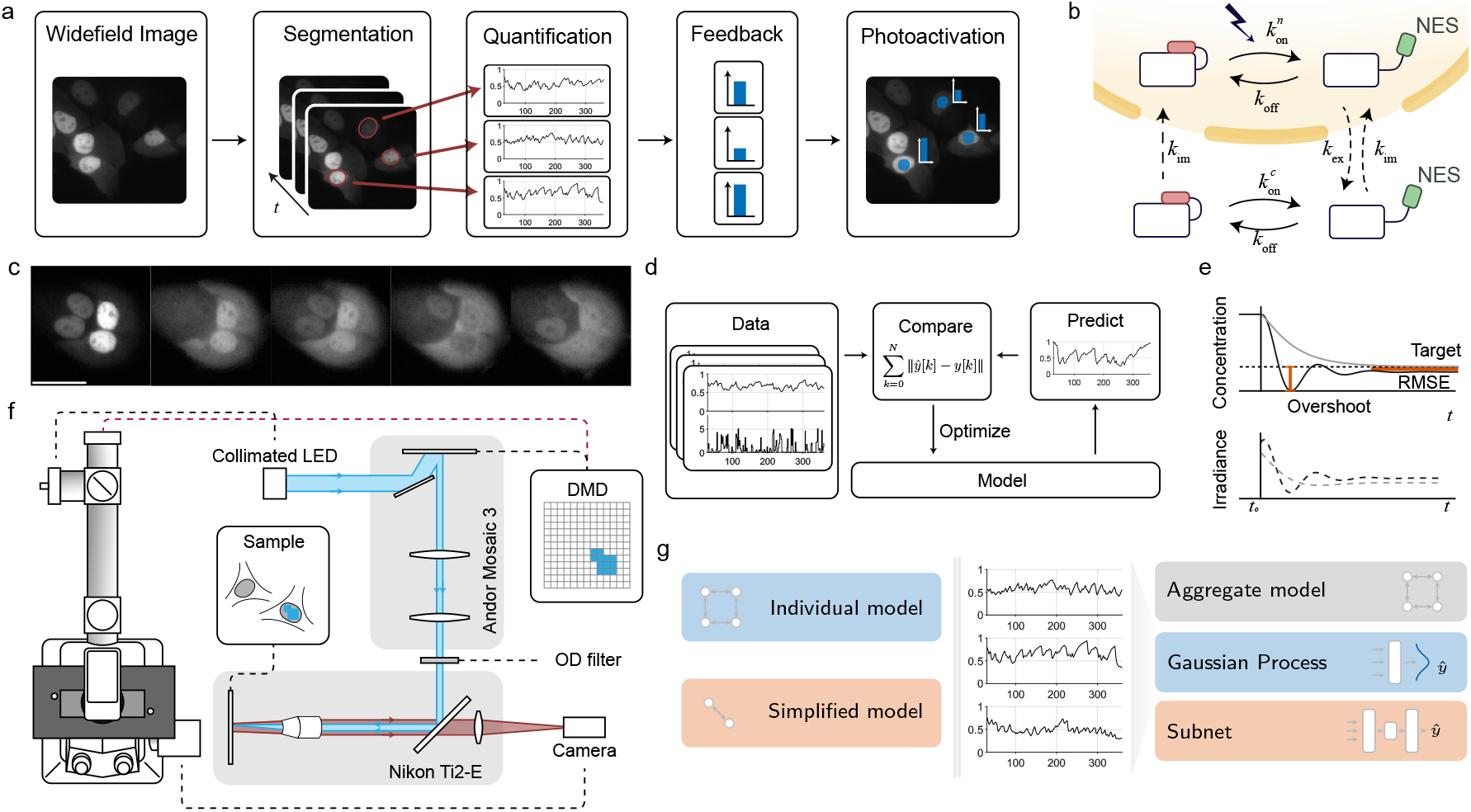
Outcome-driven microscopy for nucleocytoplasmic transport. a) Timeseries data is computed by segmenting cells and computing mean intensities to derive the optogenetic activation irradiance. This irradiance is projected onto the cell with the same segmentation mask. b) Transition dynamics and rates for the LEXY system. c) Observed image sequence under optogenetic activation of LEXY in U2OS cells (10 μm scalebar). d) Model simulations are fitted to measurement data to find model parameters. e) Past measurements up to *t*_0_ are used to compute the activation irradiance for tracking. Overshoot and RMSE are typical performance metrics used for control. f) Simplified optical path, microscope (Nikon Eclipse Ti2-E) and digital micromirror device (DMD) used to project an optogenetic stimulation mask onto the sample plane. g) Parameter identification for the physics-based and black-box models is performed at two scales: for each cell individually (left) and for the aggregated dataset (right).

While control theory offers powerful methodologies to quantify biophysical parameters and steer cellular processes, its application within the complexities of mammalian cell biology remains limited. This engineering framework uses real-time measurements to dynamically adjust input to steer system behaviour. Although limited in mammalian systems, these techniques are actively being used with optogenetics to uncover the nonlinear relationship between neural circuitry and the dynamics underlying sensory, motor and cognitive signalling (3, 14, 15), as well as to activate gene expression in bacteria or yeast to steer growth dynamics (16–20). Recent technical developments focus on control for the high stochasticity of gene expression, or use machine learning to expand control to large cell populations (21–23). Despite these advancements, standard techniques for optogenetic control have yet to be benchmarked in the heterogeneous and time-varying context of mammalian cell biology.

Benchmarking modelling and control techniques in mammalian biology requires a balance between biological complexity and experimental stability. Our recent work demonstrated subcellular control over nucleocytoplasmic transport and cell migration in mammalian cells (13), driving cells to user-defined states using closed-loop feedback. We used the Light-Inducible Nuclear Export System (LEXY) to modulate the nucleocytoplasmic transport (24). This system demonstrated high repeatability and fast dynamics under low phototoxicity, making LEXY an excellent platform for benchmarking modelling and control techniques.

In this study, we use LEXY to validate control-theoretic methodologies within mammalian cell biology. U2OS-LEXY cells were placed in an outcome-driven microscopy setup to study and control nucleocytoplasmic transport under optogenetic irradiance of the nucleus (Fig. 1a). First, we apply system identification techniques to construct predictive models by minimising the root-mean-square error (RMSE) (Fig. 1d). This step includes: rate parameter estimation in a biophysical model for mechanistic understanding, and parameter estimation in Gaussian Processes (GP) and deep subspace encoders (SUBNET) models for black-box identification (25, 26). Second, we quantify the closed-loop steering accuracy by calculating the RMSE and overshoot (Fig. 1e) under Proportional-Integral (PI) control, Linear Quadratic Regulation (LQR) and Model Predictive Control (MPC).

## METHODS

This section is structured to reflect a controller synthesis problem. Here, a model consists of quantitative equations to predict measurement evolution under optogenetic perturbation. We first introduce two biophysical prediction models for rate parameter estimation, and two black-box prediction models for simulation without explicit mechanistic understanding. Second, we use the biophysical models to implement and analyse two model-free and two model-based controllers to track a reference intensity trajectory *r*. These controllers steer the normalised nuclear intensity *y* by modulating the activation irradiance *u*, aiming to minimise the tracking error *e* = *r* − *y* over time. We end with technical details of the microscopy system and sample preparation.

### Preliminaries

LEXY drives nucleocytoplasmic transport under optogenetic illumination of the nucleus by unfolding a nuclear export signal (NES), inducing transport of the LEXY protein from the nucleus to the cytosol. This transport process is made reversible by including a nuclear localisation signal (NLS) (Fig. 1b). Finally, the fluorescent mCherry marker reveals these dynamics as intensity changes over time in fluorescence microscopy (Fig. 1c). In all experiments, we restricted both our intensity measurements and optogenetic activation to the nucleus. Restricting the light input to the nucleus enabled perturbation of cells in close proximity, without unintentionally activating their neighbours.

We normalised the nuclear intensity measurements by dividing with initial nuclear intensity, both for model identification and control experiments. This initial nuclear intensity is the baseline dark-state measurement corresponding to the maximum achievable intensity. The linearity of our biophysical model suggested that this initial intensity scales the data without altering the underlying dynamics. In this application, the normalisation improves numerical consistency during optimisation and facilitates a better comparison of results across cell-to-cell variation.

We write *y* for the normalised nuclear intensity measurements and denote measurement evolution over time as *y*(*t*) . We take these measurements at discrete time instances over intervals of length *τ*, with notation *y*[*k*] = *y*(*τk*) . We denote the predicted intensity at these time instances as *ŷ*[*k*] . Similarly, we write *u*[*k*] and *u*(*t*) for the optogenetic activation irradiance in discrete and continuous time, respectively. This optogenetic activation is held constant over the time interval from *u*[*k*] to *u*[*k* + 1].

### Identification of predictive models

#### Biophysical model

We used a four-compartment model to predict temporal dynamics of protein concentration under prior biophysical knowledge (Fig. 1b) (13). This model partitions the system into nuclear and cytosolic concentrations of active and inactive species, incorporating the rate parameters explicitly (Supp. Inf. Sec. 1A). For this model, we used an affine dependence on the activation irradiance *u* to model the photo-activation rate, 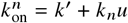 (27), where *k*^′^ is the dark-state activation rate and *k*_*n*_ the light-dependent activation rate. Using these rate kinetics then yields a differential equation for the internal concentrations 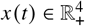 and the normalised nuclear intensity predictions *ŷ* ∈ ℝ_+,_

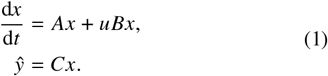

Here, the matrices *A, B* ∈ ℝ^4×4^ and *C* ∈ ℝ^1×4^ are parameterised by the kinetic rates in Figure 1b, and are explicitly defined in the Supplemental Information.

We considered a simplified two-compartment model to evaluate the predictive capabilities of our assumptions under model truncation. This model retains the bilinear structure of Eq. (1), but reduces the state vector to 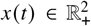 with matrices *A, B* ∈ ℝ^2×2^ and *C* ∈ ℝ^2^.

Parameters for the physics-based models were optimised under two conditions (Fig. 1g). First, a single aggregate model was fitted to data from all cells, approximating average dynamics across all cells. Second, fitting an individual model for each cell separately. This latter strategy of fitting individual models for each cell separately was also applied for the simplified model.

#### Gaussian Process and SUBNET

We considered two Machine-Learning (ML) based models aimed at enabling closed-loop control in the absence of prior system knowledge. More specifically, we use ML to learn autoregressive models, which make a prediction of the next time step based on a finite history of input-output data.

The first approach employed a Gaussian Process (GP) with a squared-exponential kernel to predict a posterior from 10 preceding measurements and inputs (Supp. Inf. Sec. 1B, Fig. S1a) (25). These GP models are especially relevant for biology because they explicitly quantify uncertainty in the prediction, thereby offering tools to deal with cell-to-cell variation and stochastic dynamics.

The second approach utilised SUBNET, an encoder-decoder recurrent neural network introduced by Beintema, Schoukens, and Tóth (26). This model architecture previously demonstrated accurate predictions in nonlinear systems by decoupling system dynamics from the measurement model. Each encoder, decoder and propagation subnetwork used 2 hidden layers of 64 features and a tanh(·) activation function, reflecting the settings suggested by Beintema, Schoukens, and Tóth (26) (Supp. Inf. Sec. 1C, Supp. Inf. Sec. 1b). This network was trained with the ADAM optimiser in Pytorch (28). Hereafter, we refer to these models as the GP model and SUBNET model.

#### Optimisation and Quantification

We optimised the model parameter vector *θ* by minimising the residual between the simulated trajectory *ŷ*[*k*] and the measurement sequence *y*[*k*] under a given input sequence *u*[*k*] of length *N*, for all model architectures:

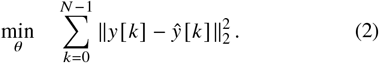

The exact prediction format for *ŷ*[*k*] is different for each model and explicitly outlined in the Supplemental Information (Supp. Inf. Sec. 1). We used the RMSE to assess the predictive performance of our models,

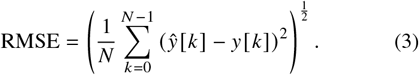

A smaller RMSE suggests a better fit of the model simulation to the trajectory measurements.

Long-term drift is rejected by closed-loop feedback in control applications. A short-horizon predictive metric is therefore more indicative of final controller performance than a full-simulation RMSE. We evaluated this short-horizon prediction accuracy for the black-box models with the *n*-step RMSE. The *n*-step RMSE quantifies the predictive performance of a model for *n* time-steps in the future,

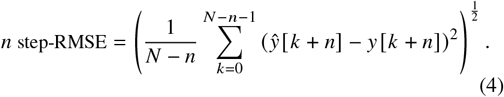

Here, *ŷ*[*k* + *n*] is the prediction made by a closed-loop simulation of the model *n* steps ahead into the future, initialised with measurement data until *y*[*k*] . For autoregressive models, this means the prediction is fed back into the model.

#### Input design for model identification

Accurate parameter identification for physical modelling requires an input signal that excites the full operating range. Practically, the activation irradiance needs to probe the full operational range of nuclear concentrations while limiting temporal dependencies to prevent biased model estimates. The irradiance should therefore prevent the system from reaching a steady-state by continuously transitioning between rise and fall kinetics. A pseudorandom sequence ensures the optimisation procedure learns system properties rather than input patterns by eliminating temporal correlations.

LEXY required a bias towards smaller activation irradiance to avoid near-constant depletion of the nucleus because the transport rate (export − import) is much quicker when most proteins are in the nucleus. To achieve this bias, we took a uniformly distributed variable *β* to induce randomness and a Bernoulli distributed variable *α* to induce sparsity, sampling

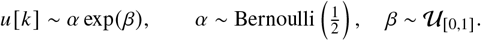

This input sequence was used to record 43 sequences, of 320 time steps each, across 8 fields of view (FOV) and over 4 coverslips, with each sequence corresponding to a different cell. Measurements and activation irradiance updates occurred at 15-second intervals, yielding 80-minute experiments of 320 measurements. The initial 180 measurements in these traces constituted the data used for fitting, and the remaining 140 measurements were used to compute validation results.

### Model-free controller design

We considered two model-independent approaches to determine the activation irradiance *u*: a proportional-integral (PI) controller with static gains and a PI controller with gains adapted to the current operating point (29, 30). These controllers combine information from the most recent error (proportional) and the past error, accumulated since the start of the experiment at *k* = 0 (integral). These controllers are commonly introduced with an additional derivative term to improve the damping-response. However, we did not use this term because it increases the sensitivity to measurement noise.

We employed gain-scheduling to improve tracking performance under the concentration-dependent transport rate of LEXY. Transport rate emerges from the product of the activation irradiance *u* and the state *x* in Eq. (1), the system therefore becomes harder to drive as the nucleus nears depletion. Consequently, the controller should favour larger activation irradiances to reach these lower nuclear intensities. To address this, our gain-scheduling strategy linearly adapted the PI gains based on the target nuclear intensity *r*, allowing a more aggressive controller at lower setpoints.

The activation irradiance is computed using the PI control law,

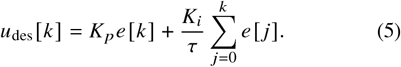

The proportional *K*_*p*_ and integral gain *K*_*i*_ are tuning parameters, designed to drive the error *e*[*k*] to 0 over time, while *τ* denotes the sampling interval.

We tuned these gains at 5 different operating points, distributed linearly over the observed operating range, resulting in 5 pairs (*K*_*p*_, *K*_*i*_). We selected the gain pair associated with the central operating point for the PI controller gains. The final gains were obtained with Matlab’s frequency response analysis and automated tuning, leveraging a linearization of the aggregate physics-based model (Supp. Inf. Sec. 2B) (31). While this approach utilised model information, the optimised gains aim to reflect achievable performance through expert manual tuning.

Further modifications were required to accommodate hardware constraints. Specifically, the laser limited the activation irradiance within the bounds *u* = min {max {*u*_des_, 0}, *u*_max_}. To mitigate integrator windup induced by this saturation, we clamped the summand in Eq. (5) at a lower and upper bound (Supp. Inf. Sec. 2B). This constraint prevented a delayed controller response by limiting error accumulation during periods of saturation.

### Model-based controller design

We considered two approaches that leverage the biophysical model to inform controller design: a linear-quadratic regulator (LQR) and model predictive control (MPC). Both approaches utilised the aggregate biophysical model internally to find the optimal control input at each time instant. The dynamics of this biophysical model in Eq. (1) are inherently nonlinear, while LQR is the optimal controller for internal state-feedback of linear systems (32, 33). Therefore, we linearised the aggregate physics-based model around a nominal operating point of *u*^∗^ = 0.1mW/cm^2^ (Supp. Inf. Sec. 2A), while maintaining the original nonlinear formulation for the MPC framework.

The resulting linearised physics-based model required a further model order reduction to enable computation of the LQR gains. To resolve this, we removed the zero-dynamics by performing a model transformation to a three-dimensional system with state *z* ∈ ℝ^3^ (Supp. Inf. Sec. 2C.1). This lower-order representation ensures the system is controllable, enabling computation of LQR gains.

To achieve tracking and disturbance rejection under parameter variation, the controller incorporates a virtual model of the reference signal and disturbance dynamics. This augmentation defines an exogenous state *w*[*k*] ∈ ℝ^6^, yielding the complete control law:

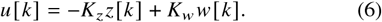

The feedback gain *K*_*z*_ ∈ ℝ^1×3^ drives the internal model state *z*[*k*] to 0 over time, while the feedforward gain *K*_*w*_ ∈ ℝ^1×6^ enforced the tracking of the reference trajectory *r*[*k*] . Together, these gains ensured that the output *y*[*k*] followed the reference signal *r*[*k*], while rejecting offsets from modelling errors and disturbances.

We computed the feedback gain *K*_*z*_ by minimising the infinite-horizon LQR cost function

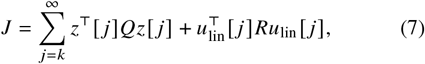

subject to the linearised closed-loop dynamics of the linearised aggregate model, setting *K*_*w*_ = 0 for stabilisation. The feedforward gain *K*_*w*_ was subsequently determined by solving the regulator equations (Supp. Inf. Sec. 2C.2).

The state penalty matrix *Q* ∈ ℝ^3×3^ and measurement penalty *R* ∈ ℝ were designed to balance controller aggressiveness. We constructed the penalty matrices using Bryson’s rule (34), setting diagonal elements inversely proportional to their maximum admissible deviations *x*_*i*,max_ and *u*_max_ for the states and input, respectively. That is, for the *i*’th diagonal element *Q*_*ii*_ = 1 / (*x*_*i*,max_)^2^ and *R* = 1 / (*u*_max_)^2^. Qualitatively, a larger *Q*_*ii*_ causes a more aggressive response to deviations during stabilisation and tracking. Given the low phototoxicity of optogenetic inputs (13), we relaxed the penalty on activation irradiance to *R* = 1 / 10^2^ while prioritizing tracking precision with *Q*_*ii*_ = 1 / 0.01^2^ for *i* ∈ {1, 2, 3} .

Implementation of the control law (6) requires access to the internal states *z*[*k*] and external states *w*[*k*], which cannot be measured directly. Therefore, we used a Kalman filter to estimate the states *z*[*k*] and *ŵ*[*k*] from nuclear intensity measurements *y*[*k*] . We outline the exact update equations and a comparison of tuning methods for the Kalman filter in the Supplementary Information (Supp. Inf. Sec. 2C.4).

MPC utilises a numerical solver to optimise the input by simulating the model over a finite horizon, minimising the deviation between the predicted and target trajectory. We again incorporated an exogenous system to account for cell-to-cell variation (Supp. Inf. 2D), mirroring the approach used in the LQR formulation.

MPC computes the optimal input sequence by solving an optimisation problem over the prediction horizon *N*_*h*_:

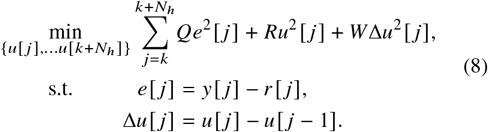

This optimisation problem is also subject to the system dynamics, detailed in the supplemental information (Supp. Inf. Sec. 2D). The problem was implemented in the `do-mpc` Python package, which leverages Casadi and the IPOPT numerical solver (35–37).

The objective function (8) seeks to minimise the tracking error *e*[*j*] while penalising the control effort *u*[*j*] and the rate of change in input signal Δ*u*[*j*] . The weighting factors *Q, R, W* ∈ ℝ_+_ were designed to balance controller aggressiveness. We again followed Bryson’s rule of defining weights as the maximum squared deviation, setting *Q* = 1 / 0.1^2^ and *R* = 1 / 10^2^. The additional derivative penalty *W* = 1 ensured smoothness and improved stability at steady-state.

This model-based MPC formulation in (8) again relies on estimates for the initial states *x*_0_ and *w*_0_ at timestep *k*. Due to the nonlinear architecture, we estimated these initial states using an extended Kalman filter (EKF) applied to a discretisation of the LEXY model (Supp. Inf. Sec. 2D.1) (38). The supplemental information also presents a comparison of tuning methods to guide future work.

#### Controller validation experiments

We evaluated the tracking efficacy of the model-free and the model-based controllers using both piecewise-constant and time-varying reference signals. The piecewise reference signal assessed steady-state tracking using an intensity sequence at *r*[*k*] = 0.7, 0.85, 0.5, 0.6, with each setpoint held constant for 20 minutes. The time-varying signal evaluated dynamic tracking and utilised a sinusoidal signal with an intensity range from 0.5 to 0.83. The frequency of this sinusoid increased linearly from 0.2 mHz to 2 mHz over 80 minutes.

### Experimental setup

For segmentation, we used the Segment Anything Model (39) in combination with a temporal filtering pipeline to extract nuclear intensity over time. Details of this image processing pipeline and the associated smart microscopy platform are detailed in our previous work (13).

For imaging, we used a Nikon Ti2-E dual-turret inverted microscope with a 40× CFI Plan Fluor NA 1.3 oil immersion objective (MRH01401, Nikon, Tokyo, Japan), placed in an incubator (Tokai-Hit, Bala Cynwyd, US) with a Sona-6 Extreme sCMOS camera (Andor, 4BV6U, Oxford Instruments, High Wycombe, UK). For fluorescence, the sample was illuminated with a CoolLED pE-4000 (CoolLED Ltd., Andover, UK) light source at 550 nm, with ET 514 nm Laser Bandpass (49905, Chroma Technology Corp., Vermont, US) and ET mCherry (49008, Chroma) filter cubes in the lower filter turret.

For activation irradiance, a 470 nm pE-800 CoolLED light source was combined with a Mosaic 3 digital mirror device (DMD) (Andor, Oxford Instruments). To reduce the activation irradiance at the sample plane, a neutral density filter with an optical density of 2.0 (NE220B, ThorLabs, Newton, US) was placed in the light path. Furthermore, a 10/90 beamsplitter (BSN10R, ThorLabs) was situated in the upper filter turret of the microscope. A disk of radius 3.6 μm was projected at the centroid of the nucleus segmentation mask using the DMD, which led to the most consistent results.

The U2OS-LEXY cells were obtained following the protocol in our previous work (13). For imaging, cells were seeded onto 25 mm coverslips, 12-24 hours before imaging, with 2 μg/mL doxycycline-hyclate (ab141091, Abcam, UK) added to induce LEXY expression. The coverslips were ultimately mounted in Attofluor cell chambers (A7816, Invitrogen).

## RESULTS AND DISCUSSION

First, we evaluate the identification performance across cells for the biophysical model and the black-box models. We then assess the model-free and model-based control architectures on both piecewise-constant and time-varying reference signals.

### Biophysical model identification

All physics-based models successfully capture the observed dynamics (Fig. 2). An example of these dynamics is presented in Figure 2a, which is one of the nuclear intensity sequences used for identification of the models. The resulting models closely replicate this measurement sequence during simulation (Fig. 2b). They also achieve comparable validation RMSE scores, although the individual models outperform the aggregate and simplified variants (Fig. 2c).

**Figure 2.**
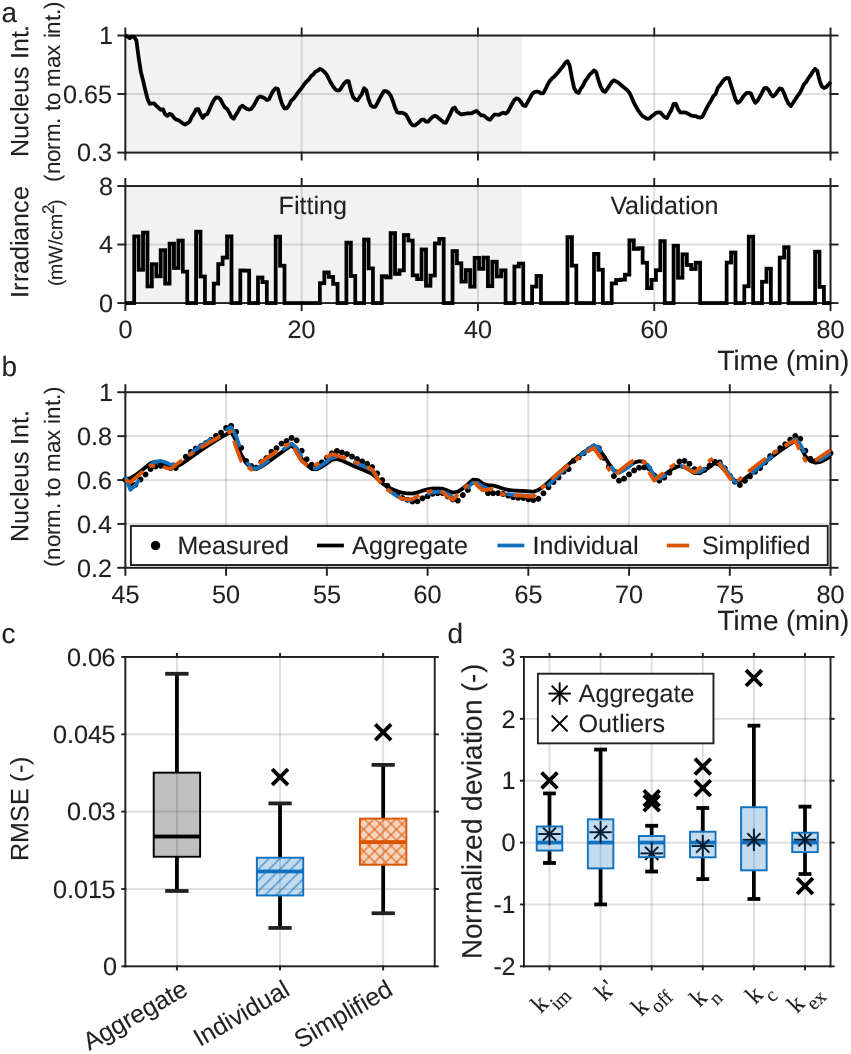
Physics-based model identification for nucleocytoplasmic transport. a) Example trace of nuclear intensity under optogenetic perturbation over time. The data is split into a training dataset and a validation dataset at measurement 180. b) Comparison of simulated and measured nuclear intensity for the aggregate, individual and simplified models. c) Simulation RMSE for the aggregate, individual and simplified models. The presented median and quartiles are computed for the RMSE of each individual nuclear intensity trace (*n* = 43). d) Normalised deviation of parameters in the individual models and the aggregate model (*n* = 43). Parameter distributions are normalised against their median.

Optogenetic perturbation provides the data richness necessary to capture biological heterogeneity without overfitting. The individual models capitalise on this rich data, outperforming the aggregate model on unseen validation data and yielding parameter distributions that cluster around the aggregate estimate (Figs. 2c, 2d). This parameter estimation is possible because persistent excitation generates input-output dynamics that are more informative than step or impulse responses (40).

High simulation accuracy suggests valid assumptions on uniform optogenetic activation and concentration-dependent intensity. These assumptions lead to tight parameter distributions for the individual models, localising parameter variability largely to the cytosolic light-dependent activation rate *k*_*c*_ and in the dark-state activation rate *k*^′^ (Fig. 2d). The observed variation in *k*_*c*_ likely stems from scattering effects, whereas the distribution of *k*^′^ currently lacks a mechanistic explanation. While our 2D application tolerates these scattering effects, additional scattering and focal plane variations will limit model accuracy in thicker 3D samples. Applying these optogenetic modelling techniques to such 3D samples will therefore require advanced spatial modelling and may benefit from fluorescence correlation spectroscopy for explicit concentration measurements.

### Gaussian Process and SUBNET identification

Both Gaussian Process (GP) and SUBNET architectures capture the nonlinear dynamics of nucleocytoplasmic transport (Fig. 3). These black-box models closely reproduce short-term behaviour, but accumulate error over horizons longer than a few minutes. This accumulating error causes long-term predictions to diverge from measurements (Fig. 3b).

**Figure 3.**
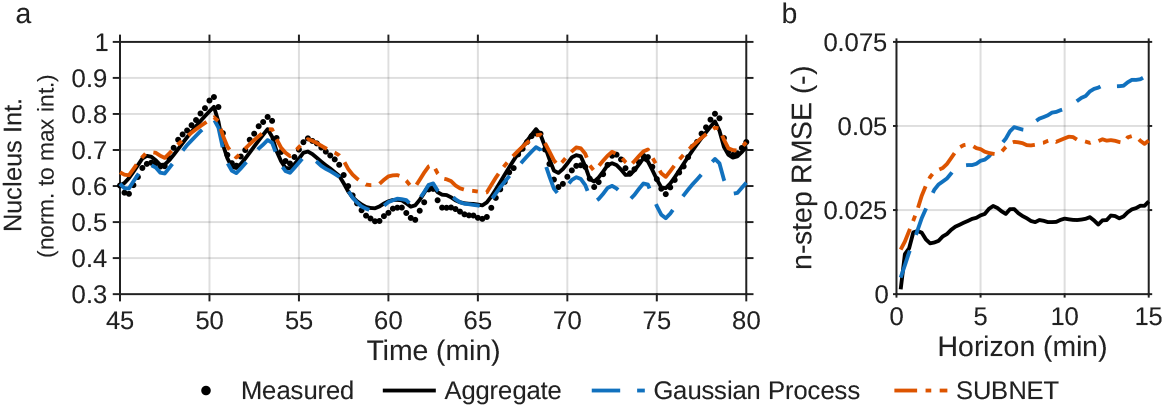
Black-box predictive model performance for nucleoctyplasmic transport. a) Simulated and measured nuclear intensity with the aggregate, Gaussian Process and SUBNET models, on the validation dataset. b) n-step RMSE of nuclear intensity for the aggregate, GP and SUBNET models over the horizon length *n*. One time-step is 15 s.

The smaller n-step RMSE for the biophysical model demonstrates that prior knowledge improves predictive performance at time-scales beyond several minutes. The biophysical model structure ensures mass conservation, directly preventing the error accumulation demonstrated by the GP and SUBNET models. The observed improvement consequently suggests that embedding physical laws into black-box models could improve their long-term predictive capabilities. For example, Physics-Informed Neural Networks can address this issue by embedding governing differential equations and physical priors directly into the loss function and structure of the network (41). These priors effectively eliminate the long-horizon integration drift while retaining model flexibility for optical and biological nonlinearities.

Despite the long-term drift, the black-box models maintain sufficient accuracy for control applications with MPC when mechanistic knowledge is lacking. This viability relies on the predictive accuracy over the initial 2.5-minute (10-timestep) horizon being only slightly worse than the aggregate model (Fig. 3b). This horizon matches typical implementations used for MPC, where the initial model accuracy dictates controller performance, rendering their long-term drift irrelevant.

### Control

All controllers demonstrate close tracking of constant setpoints across cell populations and maintain consistent behaviour across operating points (Fig. 4). The uniform behaviour leads to narrow standard deviations and similar quantiles in the traces. Similarly, the RMSE is small, and overshoot consistently remains below 10% of the setpoint (Figs. 4e and 4f). However, a direct comparison of this overshoot across controllers ignores the tuning of that specific controller. For instance, LQR and MPC controllers demonstrate larger over-shoots than the PI and GS controllers (Fig. 4f), but compensate with faster response times and lower total tracking RMSE (Figs. 4a-4e). While all controllers can match these fast response times if their gains are increased, such tuning typically increases the activation irradiance used and amplifies their sensitivity to measurement noise. Biologically, this higher activation irradiance due to noise increases the phototoxic dose on the cells.

**Figure 4.**
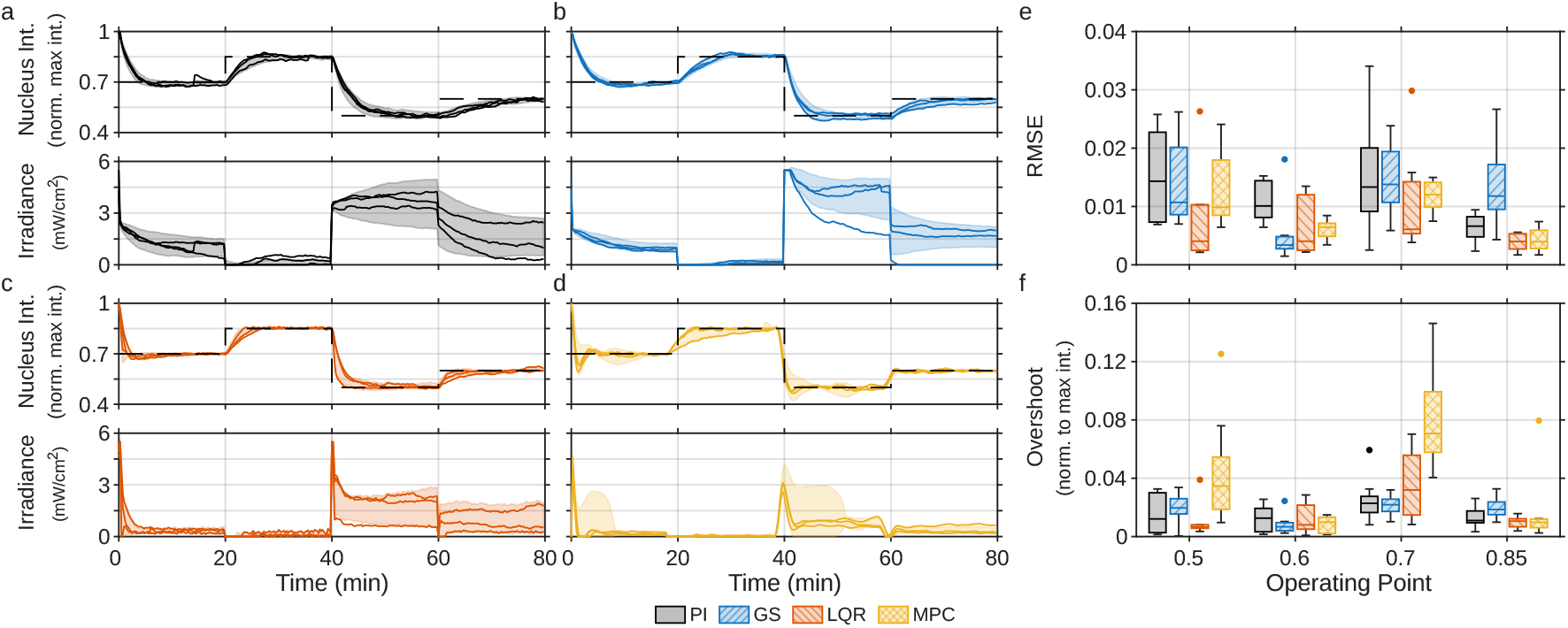
Closed-loop nuclear intensity response for PI (a, *n* = 16), GS (b, *n* = 12), LQR (c, *n* = 8) and MPC control (d, *n* = 9), tracking a piecewise constant setpoint. The shading is the pointwise mean ± standard deviation over all experiments with that controller. The continuous lines are the traces closest to 25%, 50% and 75% quantiles in total accumulated absolute error. e) RMSE over the last 10-minute segment at each constant setpoint, for each experiment. f) Absolute overshoot when changing setpoint.

LQR and MPC demonstrate robustness against variation in optogenetic sensitivity. We performed these experiments with a different cell culture than that used for the identification data and the PI controllers. Despite this new culture exhibiting higher optogenetic sensitivity, the controllers maintained close regulation (Figs. 4c-4d). This close regulation with models based on a separate biological culture suggests the LQR and MCP frameworks are robust across cell culture variability.

LQR shows the highest precision across operating points due to its inherent error handling and a fortunate placement of the linearization point. This linearization point (*u*^∗^ = 0.1 mW/cm^2^) corresponds to a nearly full nucleus, making the underlying linear model well-suited to sensitive cells. Since both LQR and MPC are theoretically optimal, incorporating cell-to-cell variation (Fig. 2d) into the MPC framework should enable performance that matches or exceeds that of LQR.

Dynamic, time-varying reference trajectories demonstrate the advantage of predictive control architectures (LQR and MPC) over the purely reactive PI and GS controllers (Fig. 5). The PI and GS controllers introduce a phase delay that worsens as the reference trajectory frequency increases. In contrast, the predictive controllers preemptively adapt the activation irradiance by leveraging the integrated exogenous model (LQR) or future trajectory (MPC). This capability largely eliminates the delay, leading to a reduced RMSE over the full 80-minute trajectory (Fig. 5e).

**Figure 5.**
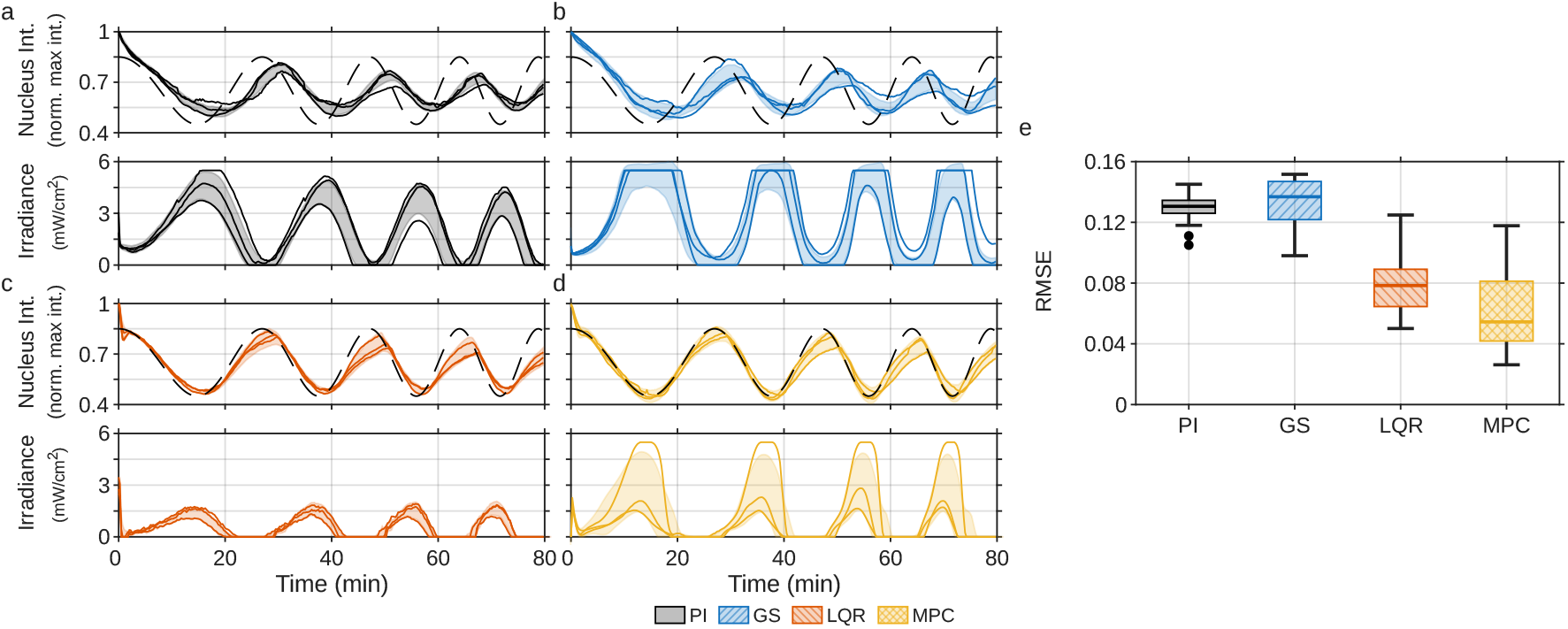
Closed-loop nuclear intensity response for PI (a, *n* = 17), GS (b, *n* = 17), LQR (c, *n* = 10) and MPC control (d, *n* = 18), tracking a time-varying setpoint. The shading is the pointwise mean ± standard deviation over all experiments with that controller. The continuous lines are the traces closest to 25%, 50% and 75% quantiles in total accumulated absolute error. e) Distribution of total accumulated RMSE for each experiment.

Notably, the GS controller exhibits a larger RMSE median and variance in the time-varying application (Fig. 5e). This originates from the more conservative tuning at high nuclear intensities. The GS controller therefore reacts slower at the peaks of the reference signal than the PI controller introducing a phase shift. However, this adaptive tuning also enabled faster response times for the piecewise constant reference signal near nucleus depletion (Figs. 4a and 4b).

## CONCLUSION

In this study, we have demonstrated that standard control-theoretic methodologies are readily applicable to mammalian cell biology by benchmarking the efficacy of system identification and control techniques for nucleocytoplasmic transport. Our results surpassed the model prediction and controller performance from our previous work by following a structured approach for the control problem (13). These models and controllers successfully capture the biological heterogeneity and enable tight control over nucleocytoplasmic transport.

Dynamic optogenetic perturbations provide sufficient data richness to achieve single-cell model identification from individual traces. These dynamic perturbations help overcome challenges typically encountered within systems biology, such as cell-to-cell variability and incomplete state observation. Additionally, they enable the automated derivation of a biological model before a control experiment if heterogeneity is otherwise difficult to overcome. We detailed an input design approach in our methods section to maximise the information content in these measurements. In our application, 180 measurements sufficed to identify an accurate model, though the necessary sequence length is dependent on model complexity and the number of free parameters. When explicit biophysical models are difficult to formulate, and the initial experiment length is limited, Gaussian Processes offer good predictions and explicit uncertainty estimates. Alternatively, if such an initial pre-experiment is not possible, for example due to phototoxicity, emerging direct data-driven control paradigms offer a promising solution (42).

There are two main limitations in applying the presented techniques to other biological systems. First, our cells had a high LEXY expression level, rendering the stochastics in reaction kinetics negligable. Conversely, these stochastic effects dominate when working with low protein counts. These stochastics invalidate the deterministic assumptions during biophysical model identification, and cause stochastic variation in trajectory tracking that cannot be overcome. Second, these techniques have computational limitations. For problems that scale towards thousands of cells in large fields of view, the computational overhead of the nonlinear solver in MPC becomes intractible. For such problems, one can leverage Deep MPC paradigms (21), or approximate the optimal control policy by extracting an explicit MPC expression. These techniques speed up the optimisation routine by enabling back-propagation through the model (Deep MPC) or by compressing the optimisation problem (explicit MPC) to locally convex problems, ensuring scalability across vast populations.

Our findings translate into practical guidelines to enable control with optogenetics. For basic tracking tasks, such as reaching a concentration setpoint with LEXY, basic PI controllers provide sufficient performance. This design requires minimal characterisation, and practical guidelines such as Ziegler-Nichols tuning are widely available (29). A gain scheduling controller can help overcome the inherent bilinearity of the system when better performance over the entire operating range is needed, or extreme setpoints need to be reached. The scheduled variable should be closely correlated with the inactive optogenetic protein concentration, because of the product-term interaction between optogenetic irradiance and protein concentration. This variable corresponded to the nuclear concentration setpoint in our application.

MPC offers the most capable framework for more complex multi-variable systems, time-varying reference tracking, or strict constraint handling. While classical techniques like LQR required explicit domain knowledge to mitigate the inherent controllability induced by mass conservation, modern MPC is flexible to implement. A variety of MPC software is available that integrates directly with data-driven black-box models, enabling control without the overhead of domain knowledge or state-estimation filters. However, as tracking requirements become more strict, the underlying cell-to-cell variation becomes more important, necessitating robust or stochastic frameworks to preserve consistent performance across population heterogeneity.

We expect that this study will enhance future outcome-driven experiments. Each of these studies will have different goals, requirements and applications, for which the evaluated methodologies in this study function as a starting point. The ability of predictive models to capture cell-to-cell variability, and the ability of our controllers to adapt to these variations, enables strong predictive and tracking performance across cellular populations. In doing so, our framework establishes a foundation for outcome-driven microscopy to uncover new biological insights with optogenetics.

## AUTHOR CONTRIBUTIONS

M.T.P.L. and C.S.S. designed the study and wrote the manuscript. M.T.P.L., A.R. and J.B.P. executed experiments. M.T.P.L. analysed data. M.T.P.L., A.R. and I.S. wrote software. M.T.P.L. prepared figures. L.C.K. and C.S.S. both contributed funding and supervised the study, with S.S. providing additional supervision.

## ACKNOWLEDGMENTS

We thank Vincent Hellebrekers (Utrecht University) for the segmentation pipeline, Florian Berger (Utrecht University) for the conception of the research line, Wilco Nijenhuis (Utrecht University) for cell cloning and Alessandro Sartori (University of Zurich) for the U2OS Flp-In T-Rex cell line. This work was supported by the Netherlands Organization for Scientific Research (NWO) Gravitation programme IMAGINE! (project number 24.005.009).

## DATA AVAILABILITY

All code and data underlying the results presented in this paper are available at [doi].

