## Supplemental Methods for "System Identification and Control for Optogenetics in Mammalian Nucleocytoplasmic Transport"


This document contains clarifying and supporting information on the implementation of methods discussed in the main article.

**Notation** We first introduce notation for continuous-time and discrete-time dynamics. The dynamics under an optogenetic activation $u$ are dictated by the map $f_\bullet(x,u)$, representing either the derivative of the continuous-time state or the state at the next discrete-time step. In this map the subscript "$\bullet$" indicates the model variant: full/nominal model ($n$), simplified model ($s$), Gaussian Process (gp) and SUBNET (sub). These models relate the concentration $x(t)$ to normalised intensity measurements $y(t)$ via the observation map $h_\bullet$. The free optimisation parameters $\theta_\bullet$ parameterise the models. In discrete-time, we use the time index $k$ to define evolution over time intervals of length $\tau$. The discrete-time evolution for the state $x$ is then denoted $x(k\tau) = x[k]$, and similarly defined for $u$ and $y$. For the autoregressive models, we define the regressive vectors $\chi[k] = [\ u[k], u[k-1], \cdots, u[k-n_p-1], y[k], y[k-1], \cdots y[k-n_p-1]]^\top \in \mathbb{R}^{2n_p}$, where the regressor size $n_p$ is defined in context.

### 1. IDENTIFICATION

#### A. Biophysical models

We formulate the differential equations for nucleocytoplasmic transport using the four-compartment model for nucleic and cytosolic concentrations, shown in Figure 1b of the main text. The model simplifies diffusion dynamics, volumetric effects and nucleopore limitations to simple rate parameters $k_\bullet$. Our affine assumption on the optogenetic activation irradiance $u$ results in a light-dependent activation rate $k_{\text{on}}^n = k' + k_n u$ [1], with $k'$ a base dark-state activation and $k_n$ the light-dependent rate. Additionally, we account for residual activation in the cytosol $k_{\text{on}}^c = k' + k_c u$, capturing scattering effects. Altogether, this leads to a differential equation for the active ($a$) and dark-state ($d$) nuclear ($n$) and cytolic ($c$) concentrations, denoted with $n_a$, $c_a$, $n_d$ and $c_d$,

$$\begin{bmatrix} \dot n_a \\ \dot n_d \\ \dot c_a \\ \dot c_d \end{bmatrix} = \underbrace{\overbrace{\begin{bmatrix} -k_{\text{off}} - k_{\text{ex}} & k' & k_{\text{im}} & 0 \\ k_{\text{off}} & -k' & 0 & k_{\text{im}} \\ k_{\text{ex}} & 0 & -k_{\text{off}} - k_{\text{im}} & k' \\ 0 & 0 & k_{\text{off}} & -k' - k_{\text{im}} \end{bmatrix}}^{A_n} \overbrace{\begin{bmatrix} n_a \\ n_d \\ c_a \\ c_d \end{bmatrix}}^{x} + u \overbrace{\begin{bmatrix} 0 & k_n & 0 & 0 \\ 0 & -k_n & 0 & 0 \\ 0 & 0 & 0 & k_c \\ 0 & 0 & 0 & -k_c \end{bmatrix}}^{B_n} \begin{bmatrix} n_a \\ n_d \\ c_a \\ c_d \end{bmatrix}}_{f_n(x,u;\theta_n)}, \tag{S1}$$

$$\hat y = \underbrace{\overbrace{\begin{bmatrix} 1 & 1 & 0 & 0 \end{bmatrix}}^{C} x}_{h_n(x)}.$$

Here, $\hat y$ denotes the predicted normalised intensity of the nucleus. Figure 1b of the main text defines all rates $k_\bullet$. The matrices $A_n$, $B_n$ and $C$ in Eq. (S1) will be used repeatedly throughout the document, while keeping notation consistent.

Note that scaling $x$ and inversely scaling $h_n$ does not change the predicted measurements $\hat y$. This symmetry indicates that measurement data can not uniquely determine the rate parameters $k_\bullet$. Therefore, we fix

$h_n(x) = [1\ 1\ 0\ 0]\, x$, leading to a unique set of $k_\bullet$. By ensuring that these parameters are unique, we made the system structurally identifiable [2].

Experiments indicated that the sensitivity of the optogenetic switch dominates the system dynamics, with transport occurring at faster timescales. Hence, we consider a simplified 2-compartment model in which optogenetic activation acts directly on import/export rates. This simplification leads to a state-space description analogous to the four-compartment model in Eq. (S1):

$$\begin{aligned} \begin{bmatrix} \dot{n} \\ \dot{c} \end{bmatrix} &= \underbrace{\begin{bmatrix} -k_{\text{ex}} & k_{\text{im}} \\ k_{\text{ex}} & -k_{\text{im}} \end{bmatrix} \begin{bmatrix} n \\ c \end{bmatrix} + u \begin{bmatrix} -k_p & k_c \\ k_p & -k_c \end{bmatrix} \begin{bmatrix} n \\ c \end{bmatrix}}_{f_s(x,u;\theta_s)}, \\ \hat{y} &= \underbrace{\begin{bmatrix} 1 & 0 \end{bmatrix}}_{h_s(x)} x \,. \end{aligned} \tag{S2}$$

The state $x = [n\ c]^\top$ describes the total concentration of the construct in the cytosol ($c$) and nucleus ($n$), respectively. Here, the same considerations for scaling and structural identifiability apply, fixing $h_s(x)$ to the expression in Eq. (S2).

We estimate the rate parameters $\theta_\bullet$ and initial system state $x(t_0)$ by solving a constrained nonlinear least-squares optimisation problem for both the four-compartment and two-compartment models. This optimisation problem minimises the squared residual between the predicted intensity $\hat{y}$ and the measurements $y$,

$$\begin{aligned} &\min_{\theta_\bullet, x(t_0)} \sum_{k=0}^{N-1} \|\hat{y}[k] - y[k]\|_2^2 + \lambda \|\theta_\bullet\|_2^2 \\ &\text{s.t.} \quad \hat{y}[k] = h_\bullet \left( \int_{t_0}^{t_0 + k\tau} f_\bullet\left(x(t), u(t); \theta_\bullet\right) \,\mathrm{d}t \right). \end{aligned} \tag{S3}$$

We incorporate Tikhonov regularisation on the rate parameters with weight $\lambda = 0.1$ to mitigate overfitting. The optimisation problem is solved using gradient descent in Matlab's System Identification Toolbox [3]

### B. Gaussian Process

We model the black-box system dynamics using Gaussian Process (GP) in a Nonlinear Autoregressive with Exogenous Inputs (NLARX) framework. In this specific implementation, we use Matlab's GP formulation, combining a linear prediction with the GP prediction to approximate the nonlinearities (Fig. S1a):

$$\hat{y}[k+1] = f_{\text{gp}}(\chi[k]; \theta_{\text{gp}}) := y_0 + \chi^\top[k] L + G(\chi[k]; \theta_{\text{gp}}). \tag{S4}$$

Here, the next-step prediction $\hat{y}[k+1]$ represents the posterior mean derived from a linear offset $y_0 \in \mathbb{R}$, linear projection weight $L \in \mathbb{R}^{2n_p}$ and the GP kernel correction $G(\chi[k]; \theta_{\text{gp}})$. To compute this GP correction, the training data is compiled into matrices $X \in \mathbb{R}^{2n_p \times N}$ and $Y \in \mathbb{R}^{N \times 1}$, concatenating all input-output regressor pairs $(\chi[k], y[k])$.

We use the exponential kernel function for the GP correction:

$$k(\chi[k]; \chi_i; \theta_{\text{gp}}) = \sigma_f^2 \exp\left( -\frac{1}{2} \frac{(\chi[k] - \chi_i)^\top (\chi[k] - \chi_i)}{\sigma_t^2} \right),$$

where the parameter vector $\theta_{\text{gp}}$ collects the signal variance $\sigma_f^2$, the characteristic time scale $\sigma_l^2$, and the measurement noise variance $\sigma_n^2$. This kernel function then defines elements of the full training covariance matrix $K(X, X) \in \mathbb{R}^{N \times N}$ with $K(X, X)_{i,j} = k(\chi_i, \chi_j)$. Similarly, for the cross-covariance $K(\chi[k], X) \in \mathbb{R}^{1 \times N}$ with $K(\chi[k], X)_j = k(\chi[k], \chi_j)$. Finally, the GP-based output prediction is computed

$$G(\chi[k]; \theta_{\text{GP}}) = K(\chi[k], X)(K(X, X) + \sigma_n^2 I)^{-1} Y. \tag{S5}$$

Autoregressive models can optimise directly against the prediction error rather than minimising the long-term simulation error. This approach is often preferred when dealing with process noise and structural modelling errors [4]. However, physics-based models cannot utilise this approach, because their predictions depend on unobservable internal states. Thus, we minimise the prediction error to find the parameters for the linear bias $L, y_0$ and covariance parameters $\sigma_f$ and $\sigma_t$ of the GP

$$\begin{aligned} &\min_{\theta_{\text{gp}}} \sum_{k=n_p}^{N-1} \|y[k] - \hat{y}[k]\|_2^2 \\ &\text{s.t.} \quad \hat{y}[k] = y_0 + \chi^\top[k] L + G(\chi[k], \theta_{\text{gp}}) \qquad k = n_p, n_p + 1, \ldots, N-1 \end{aligned} \tag{S6}$$

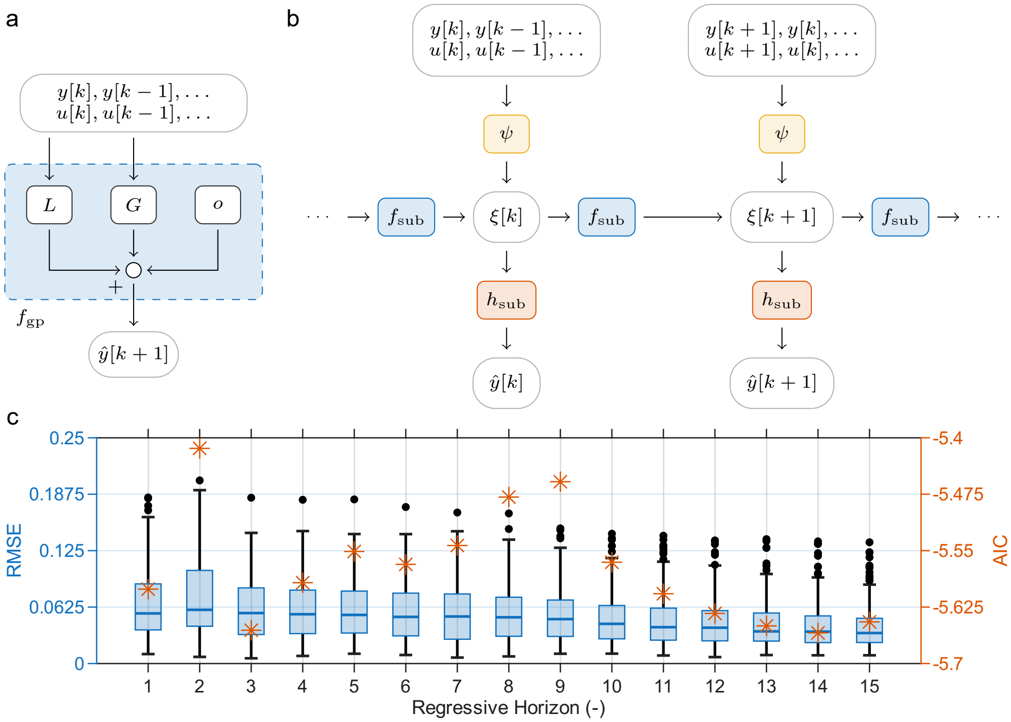


**Fig. S1.** a) The Gaussian Process $G$, a linear component $L$ and an offset $o$ makes a prediction $\hat{y}[k+1]$ from past-input output data with the transition map $f_{\text{gp}}$. b) SubNet is an autoencoder with a Recursive Neural Network propagated by $f_{\text{sub}}$, where past input-output data is lifted into a latent space using $\psi$ and measurement estimates are recovered using $h_{\text{sub}}$. c) Simulation RMSE over 250 simulations, and Akaike's Information Criterion (AIC) for Gaussian Process predictions at different past horizons.

To select the past horizon length $n_p$, we investigated the simulation Root Mean Square Error (RMSE) and Akaike's Information Criterion (AIC) across different autoregressive horizons on the validation dataset (Fig. S1c). The AIC is defined as [5],

$$\mathrm{AIC} = \log\left(\frac{1}{N-n_p}\sum_{k=n_p}^{N-1}(y[k]-\hat{y}[k])^2\right) + \frac{2N_p}{N}, \tag{S7}$$

where $N_p$ is the number of parameters in the model, and $N$ the number of predictions used for the estimate. The AIC quantifies the tradeoff between predictive accuracy and the cost of additional parameters, minimising this criterion should therefore select for the most parsimonious model.

Figure S1c suggests a model order of either 3 or 14 with diminishing returns for the RMSE and AIC. Because we are not limited computationally and want to gauge optimal performance, we used a model horizon of $n_p = 14$.

#### C. Subnet

The encoder/decoder RNN model for system identification reflects that of a state-space model with internal states (Fig. S1b). Inspired by the four-compartment physical model, we select a latent space dimension of 4 for the internal state vector $\xi[k] \in \mathbb{R}^4$. The network includes an encoder $\psi : \mathbb{R}^{14} \to \mathbb{R}^4$ to initialise the internal state $\xi[k]$ from historical data, a latent-space propagator $f_{\text{sub}} : \mathbb{R}^4 \times \mathbb{R} \to \mathbb{R}^4$ to drive transport dynamics, and a decoder $h_{\text{sub}} : \mathbb{R}^4 \to \mathbb{R}$ to recover the measurement predictions:

$$\begin{aligned} \xi[k] &= \psi(\chi[k], \theta_{\text{sub}}), \\ \xi[k+1] &= f_{\text{sub}}\left(\xi[k], u[k]; \theta_{\text{sub}}\right), \\ \hat{y}[k] &= h_{\text{sub}}\left(\xi[k]; \theta_{\text{sub}}\right). \end{aligned} \tag{S8}$$

Following suggestions by Beintema et al. [6], each of the networks $\psi$, $f_{\text{sub}}$ and $h_{\text{sub}}$ is parameterised as a multilayer perceptron with 2 hidden layers of 64 features each and hyperbolic tangent ($\tanh(\cdot)$) activation functions. The encoder network $\psi$ uses past input-output elements for $\chi[k]$ to determine the internal state $\xi[k]$. We again used the past 14 measurements to match the horizon used for the GP formulation.

The network parameters $\theta_{\text{sub}}$ were trained by minimising the multi-step prediction error over a prediction horizon of $T = 20$:

$$\begin{aligned} \min_{\theta_{\text{sub}}} \; & \sum_{k=n+1}^{N_h-T+1} \sum_{i=0}^{T-1} \|y[k+i] - \hat{y}[k+i]\|_2^2 \\ \text{s.t.} \; & \xi[k+i+1] = f_{\text{sub}}(\xi[k+i], u[k+i]), \qquad i = 0, \ldots, T-1 \\ & y[k+i] = h_{\text{sub}}(\xi[k+i]), \qquad i = 0, \ldots, T-1 \\ & \xi[k] = \psi(u[k], \ldots, y[k], \ldots) \end{aligned} \tag{S9}$$

We implemented and optimised this network in the `deepSI` Python package, using `PyTorch` with the ADAM optimiser and a batch size of 256 for 5000 iterations (learning rate 0.001, momentum parameters $\beta = (0.9, 0.99)$) [7, 8]. Optimisation took around 5 minutes on an Intel i9-13900H CPU.

### 2. CONTROL

#### A. Discretisation and linearisation

To analyse and design linear control methods that act in discrete time, we first discretise and then linearise the full four-compartment model from Eq. (S1) around an equilibrium point $(x^*, u^*)$. To discretise the model in Eq. (S1) we can compute an analytical solution under constant activation irradiance $u[k]$ over a time interval $\tau$,

$$x[k+1] = F(x[k], u[k]) := \int_{k\tau}^{(k+1)\tau} f_n(x(t), u[k]) \; \mathrm{d}t = e^{(A_n + u[k]B_n)\tau} x[k] \tag{S10}$$

This solution is a matrix exponential with matrices $A_n$ and $B_n$, referring to the matrices in the underlying dynamics in Eq. (S1).

Next, we linearise this matrix exponential around an operating point indexed by $i$ to obtain a linear model in standard state-space notation with deviation variables $x_{\text{lin},i} := x(t) - x_i^*$ and $u_{\text{lin},i} := u(t) - u_i^*$ (Fig. S2a),

$$x_{\text{lin},i}[k+1] = \underbrace{\left.\frac{\partial F}{\partial x}\right|_{x_i^*, u_i^*}}_{A_{l,i}} x_{\text{lin},i}[k] + \underbrace{\left.\frac{\partial F}{\partial u}\right|_{x_i^*, u_i^*}}_{B_{l,i}} u_{\text{lin},i}[k] \tag{S11}$$

In practice, we numerically estimate this derivative using a central difference method because the derivative of this matrix exponential is nontrivial.

### B. Proportional-Integral controller design

We tuned our PI and GS controllers across a range of target operating points. The baseline PI controller uses a fixed gain pair $(K_p, K_I)$, whereas the GS controller switches the gain pair based on the active operating region. Each of these gains was determined optimally at operating points $y_i^* \in \{1, 0.875, 0.75, 0.625, 0.5\}$. We numerically inverted the operating point estimate from the model to determine the underlying state $(x_i^*, u_i^*)$, required for tuning these gains,

$$x_i^*(u_i^*) = \lim_{t\to\infty} e^{(A_n+u_i^* B_n)t} x_0, \qquad y_i^*(u_i^*) = Cx_i^*(u_i^*), \tag{S12}$$

where the initial state $x_0$ was determined during the identification procedure. This matrix exponential is again the solution of the differential equation in Eq. (S1) under constant activation irradiance $u_i^*$. In practice, we computed this limit by intersecting the space of constant protein count at the initial condition $x_0$ with the null space of the dynamics (i.e. where $A_n + u_i^* B_n = 0$) with:

$$\text{Null}\left(A_n + u_i^* B_n\right) \cap \{x \ : \ \|x\|_1 = \|x_0\|_1\}.$$

Controller design was guided by frequency-domain loop-shaping analysis [9, 10]. The main goals in controller design is to obtain reference tracking to some reference trajectory, and to reject disturbance effects on our measurements. Therefore we focus on the closed-loop transfer function $T(\omega; y_i^*)$, mapping the reference input $r$ to the output $y$, and the sensitivity function $S(\omega; y_i^*)$, which maps the disturbance $d$ to the output $y$ (Fig. S2b). In these transfer function we use the index $i$ to index the operating points for the linearisation used in Eq. (S11). These transfer functions describe the steady-state response of the output $y$ under a sinusoidal forcing for $r$ or $d$ at frequency $\omega$. We compute these transfer functions by evaluating a DTFT of the input-output dynamics in Eq. (S11),

$$T_i(\omega; y_i^*) = \frac{M_i(\omega; y_i^*)P_i(\omega)}{1 + M_i(\omega; y_i^*)P_i(\omega)}, \qquad S_i(\omega) = \frac{1}{1 + M_i(\omega; y_i^*)P_i(\omega)}.$$

Here, $M_i(\omega; y_i^*) = C(\exp(j\omega)I - A_{l,i})^{-1}B_{l,i}$ is the transfer function associated with Eq. (S11) evaluated at the equilibrium $y_i^*$, and $P_i(\omega) = K_p + K_I\tau/(\exp(j\omega) - 1)$ is the transfer function associated with the PI controller, with $j$ the complex unit.

We aimed toa achieve both tracking with an overshoot of a maximum of 5 % and noise rejection. These goals translate into a peak resonant gain of $\|T_i(\omega; y_i^*)\| \leq 1.05$ for the transfer function, and a peak gain of $\|S_i(\omega; y_i^*)\|_\infty \leq 1$ for the sensitivity function. We initially computed controller gains with the Matlab `pidtune` command [11], and subsequently refined them manually to achieve our targets (Fig. S2d). We present the final tuned transfer functions across all operating points for the aggregate and individual models in Figure S2c, with the final gains stated explicitly in S1.

**Table S1.** Controller gains used for Proportional-Integral control and Gain Scheduling at measured normalised intensity operating points $y$.

| | PI | $y = 1$ | $y = 0.875$ | $y = 0.75$ | $y = 0.625$ | $y = 0.5$ |
|---|---|---|---|---|---|---|
| $K_p$ | 8 | 2 | 4 | 6 | 8 | 15 |
| $K_I$ | 0.05 | 0.0075 | 0.02 | 0.03 | 0.05 | 0.075 |

The activation irradiance was bounded by hardware constraints of the LED system, such that $u \in [0, u_{\max}]$. These bounds can cause the integrator state of the PI controller to accumulate error when the requested input cannot be realised. For example, if the nucleus is nearly depleted (intensity $y = 0.5$), it can take a while for the nucleus to recover towards a high reference target of $r = 1$. Meanwhile, the error $e = y - r$, remains negative (requesting $u < 0$), causing the integrator to accumulate a large negative component and deteriorating controller performance. This integrator windup was avoided by conditionally clamping the integrator [9], yielding a dynamic description for the PI controller,

$$\begin{aligned} I[k] &= \min\left[\max\left(I[k-1] + \frac{1}{\tau}e[k], I_{\min}\right), I_{\max}\right] \\ u[k] &= K_p e[k] + K_I I[k]. \end{aligned}$$

This PI controller was used in our implementation with bounds $I_{\max} = 10^5$ and $I_{\min} = 0.1$, and a sampling time $\tau = 15$ s.

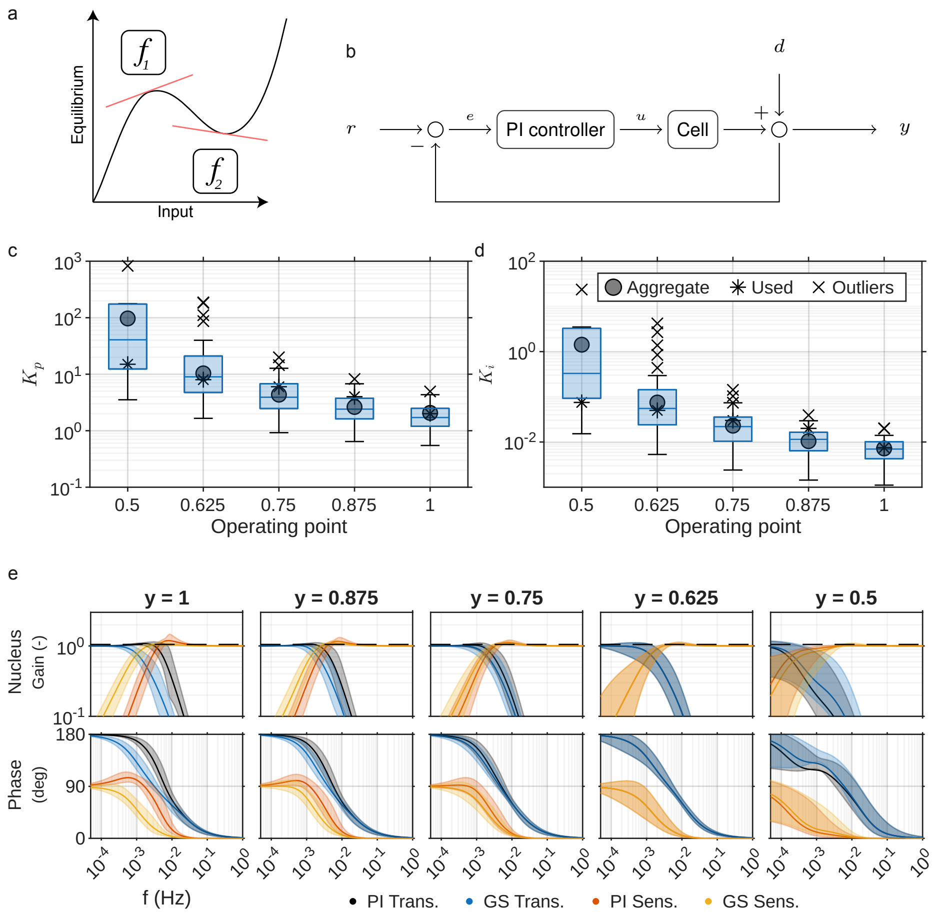


**Fig. S2.** a) Linearising a nonlinear system around an operating point. b) Diagram of signal flow, a reference $r$ and measurement $y$ are fed into a controller, which determines the optogenetic activation irradiance $u$ applied to the cell, subject to a measurement disturbance $d$. c) Distribution of Proportional gains optimised at operating points using MATLAB's autotuning tool for each cell model. d) Distribution of Integral gains. e) Closed-loop transfer (Trans.) and sensitivity (Sens.) function at designed operating points for the linearised LEXY system under PI control and gain-scheduled PI control. The continuous line represents the response of the aggregate model, whilst the shaded band indicates the mean ± standard deviation across all individual models ($n = 43$). The dashed line indicates the 5% overshoot design target.

### C. Linear Quadratic Regulator for output tracking

LQR controller is the optimal controller for the linearised, discretised dynamics in S11. To find the final gains, we follow three key steps: first we remove an uncontrollable mode from the dynamics, then we design a virtual system to capture disturbance and reference trajectory dynamics, and then we compute the final controller gains. The latter two steps are standard in control, often called the Internal Model Principle [12, 13]. The aggregate physics-informed model (Section A) demonstrated better predictive performance than black-box models, and was therefore used in this control formulation.

In what follows, we used the linearised dynamics in Eq. (S11) at a linearisation point of $u^* = 0.1$ mW/cm$^2$, with $x^*$ and $y^*$ determined with Eq. (S12). We drop the index notation $i$ for the equilibrium states and the matrices $A_l$ and $B_l$ in Eq. (S11) to indicate that we only consider a single linearisation point.

#### C.1. Mass conservation state removal

The underlying transport equations conserve total protein mass, consequently, the discretised matrix $A_l$ contains an uncontrollable eigenvalue at 1. We isolated this eigenvalue by computing the eigendecomposition of $A_l$ to construct a coordinate transform $T$. This yielded a block structure

$$A_l = T \begin{bmatrix} 1 & 0 \\ 0 & A \end{bmatrix} T^{-1}, \qquad B_l = T \begin{bmatrix} 0 \\ B \end{bmatrix},$$

where $A$ is a diagonal matrix containing the remaining eigenvalues, and $B$ is the transformed input matrix. This transformation maps the discrete-time dynamics to

$$x_{\text{lin}}[k+1] = T \begin{bmatrix} 1 & 0 \\ 0 & A \end{bmatrix} T^{-1} x_{\text{lin}}[k] + T \begin{bmatrix} 0 \\ B \end{bmatrix} u_{\text{lin}}[k].$$

We simplified these dynamics by defining a new state vector $z_c = T^{-1}x_{\text{lin}}$ to obtain a partition into the uncontrollable total protein concentration and the controllable state vector $z[k]$, yielding

$$\begin{aligned} z_c[k+1] &= \begin{bmatrix} \text{zero mode} \\ z[k+1] \end{bmatrix} = \begin{bmatrix} 1 & 0 \\ 0 & A \end{bmatrix} \begin{bmatrix} \text{zero mode} \\ z[k] \end{bmatrix} + \begin{bmatrix} 0 \\ B \end{bmatrix} u_{\text{lin}}[k]. \\ y_{\text{lin}}[k] &= \begin{bmatrix} C' & C_z \end{bmatrix} z_c[k] = CT^{-1}x_{\text{lin}}[k] \end{aligned} \tag{S13}$$

This equation reflects that the total protein concentration remains constant over time and is unaffected by the control input $u_{\text{lin}}[k]$. Consequently, we truncated this system into just the dynamics for $z[k]$ and restricted the LQR tracking problem to the dynamics of this controllable vector $z[k]$. Now, when $z[k] = 0$, $u_{\text{lin}}[k] = 0$ and $y_{\text{lin}}[k] = 0$, the linearised system is in steady state at a constant nucleocytoplasmic transport balance. This balance corresponds directly to the linearisation point $(x^*, u^*)$.

#### C.2. Exogenous system formulation

We included a virtual model of the reference signal and external disturbances to achieve reference tracking with disturbance rejection and robustness against cell-to-cell variation. This inclusion enables the controller to stabilise against future predicted setpoints and disturbances. We define this virtual model with an internal state vector $w \in \mathbb{R}^5$ as a linear discrete-time system,

$$\begin{aligned} w[k+1] &= S_{\text{lqr}} w[k], \\ e[k] &= y_{\text{lin}}[k] - C_r w[k], \end{aligned} \tag{S14}$$

where $S_{\text{lqr}} \in \mathbb{R}^{5\times5}$ represents discrete-time dynamics, designed to accommodate constant disturbances and time-varying reference signals. Furthermore, $C_r$ maps the state vector $w[k]$ to the target reference command $r[k] = C_r w[k]$, and $e[k]$ then denotes the resulting tracking error.

Integrating this virtual model with the dynamics of the controllable linearised system yields the augmented state-space representation,

$$\begin{aligned} z[k+1] &= Az[k] + Bu_{\text{lin}}[k] + B_{\text{lqr}} w[k] + \nu[k], \\ y_{\text{lin}}[k] &= C_z z[k] + C_{\text{lqr}} w[k] + \mu[k], \end{aligned} \tag{S15}$$

where $z \in \mathbb{R}^3$ is the controllable transformed state, $u_{\text{lin}}[k] := u[k] - u^*$ the input deviation and $y_{\text{lin}}[k] := y[k] - Cx^*$ the measurement deviation from the linearisation point. The virtual state vector $w[k]$ then models both process and measurement disturbances acting through $B_{\text{lqr}}$ and $C_{\text{lqr}}$, respectively. Stochastic behavior and measurement errors are captured with Gaussian noise processes $\nu[k] \sim \mathcal{N}(0, Q)$ and $\mu[k] \sim \mathcal{N}(0, R)$, governed by unknown covariance matrices $Q \in \mathbb{R}_{+}^{3\times 3}$ and $R \in \mathbb{R}_{+}$. We partition the virtual system state $w[k]$ into the individual measurement and process disturbances with $w_z[k] \in \mathbb{R}$ and $w_d[k] \in \mathbb{R}$ respectively, and into the tracking variable $w_r[k] \in \mathbb{R}^3$:

$$w[k+1] = \underbrace{\begin{bmatrix} 1 & 0 & 0 \\ 0 & 1 & 0 \\ 0 & 0 & e^{S'_{\text{lqr}}\tau} \end{bmatrix}}_{S_{\text{lqr}}} \begin{bmatrix} w_z[k] \\ w_d[k] \\ w_r[k] \end{bmatrix}.$$

Here, we assume that the state and measurement disturbances $w_z[k]$ and $w_d[k]$ evolve slowly, allowing us to model them as constant over time. Similarly, we design specific reference dynamics for $w_r[k]$ using a continuous-time generator $S'_{\text{lqr}}$, discretised over the sampling interval $\tau = 15$ s.

$S'_{\text{lqr}}$ was designed in continuous-time to achieve the simultaneous goals of piecewise-constant and oscillatory reference tracking:

$$S'_{\text{lqr}} = \text{diag}\left( \underbrace{0}_{\text{constants}}, \underbrace{\begin{bmatrix} 0 & \omega_1 \\ -\omega_1 & 0 \end{bmatrix}}_{\text{osc. at freq } \omega_1} \right).$$

A similar design can be used to deal with known non-constant disturbance dynamics. However, the modelling errors should lead to slowly changing disturbances in our application. We used a single oscillation frequency of $\omega_1 = 0.8$ mHz ($2\pi \times 8 \times 10^{-4}$ rad/s) to match the sinusoidal reference signal during controller validation. These definitions lead to explicit expressions for the final matrices in Eq. (S15),

$$B_{\text{lqr}} = \begin{bmatrix} 0 & 0 & 0 & 0 & 0 \\ 0 & 0 & 0 & 0 & 0 \\ 1 & 0 & 0 & 0 & 0 \end{bmatrix}, \qquad C_{\text{lqr}} = \begin{bmatrix} 0 & 1 & 0 & 0 & 0 \end{bmatrix}, \qquad C_r = \begin{bmatrix} 0 & 0 & 1 & 1 & 0 \end{bmatrix}.$$

We introduced a small decay term into the discrete-time transition matrix $S_{\text{lqr}}$ to obtain $S_{\text{used}} = S_{\text{lqr}} - \epsilon I$ with $\epsilon = 0.01$. This adjustment aids the stability of the closed-loop system by assuming a slow decay of any disturbances.

#### C.3. Constructing the linear quadratic regulator

With the augmented virtual state $w[k]$ and the transformed state $z[k]$ defined, we formulate the composite LQR law:

$$u[k] = -K_z z[k] + K_w w[k]. \tag{S16}$$

This law aims to drive the tracking error $e[k]$ to zero over time. The feedback gain $K_z$ acts on the state vector $z[k]$ to drive $z[k]$ to 0, whereas the feedforward gain $K_w$ rejects disturbances and shifts the equilibrium point towards the reference trajectory $r[k]$. This structure ensures that the output $Cz[k]$ tracks the reference path while rejecting offsets due to disturbances and modelling errors.

We minimised the infinite-horizon LQR cost function to compute the feedback gain $K_z \in \mathbb{R}^{1\times 3}$,

$$J = \sum_{j=k}^{\infty} z^\top[j] Q_{\text{lqr}} z[j] + u_{\text{lin}}^\top[j] R_{\text{lqr}} u_{\text{lin}}[j], \tag{S17}$$

subject to the dynamics in equations Eq. (S14) – Eq. (S16) with $K_w = 0$. The positive semidefinite matrix $Q_{\text{lqr}} \in \mathbb{R}_{+}^{3\times 3}$ and positive definite scalar $R_{\text{lqr}} \in \mathbb{R}_{>0}$ serve as tuning parameters to balance controller aggressiveness. A desciption of the used cost matrices is discussed in the main text.

We computed the remaining feedforward gain vector $K_w \in \mathbb{R}^{1\times 5}$ by solving the discrete-time regulator equations for $\Pi \in \mathbb{R}^{3\times 5}$ and $\Gamma \in \mathbb{R}^{1\times 5}$,

$$\begin{bmatrix} A & B \\ C & 0 \end{bmatrix} \begin{bmatrix} \Pi \\ \Gamma \end{bmatrix} - \begin{bmatrix} \Pi \\ 0 \end{bmatrix} S + \begin{bmatrix} B_{\text{lqr}} \\ C_{\text{lqr}} - C_r \end{bmatrix} = 0.$$

We used the Kronecker-product (denoted by $\otimes$) vectorisation identity to solve this equation directly, $\operatorname{vec}(ABC) = \left(C^\top \otimes A\right) \operatorname{vec}(B)$. This identity transforms the algebraic relation into a linear equation,

$$-\left(I \otimes \begin{bmatrix} A & B \\ C & 0 \end{bmatrix} - S^\top \otimes \begin{bmatrix} I & 0 \\ 0 & 0 \end{bmatrix}\right)^{-1} \operatorname{vec}\left(\begin{bmatrix} B_{\text{lqr}} \\ C_{\text{lqr}} - C_r \end{bmatrix}\right) = \operatorname{vec}\left(\begin{bmatrix} \Pi \\ \Gamma \end{bmatrix}\right), \tag{S18}$$

where $I_\bullet \in \mathbb{R}^{\bullet \times \bullet}$ indicate identity matrices with the indicated size. Ultimately, these solutions combine into the full feedforward gain $K_w$ in Eq. (S16) by computing

$$K_w = \Gamma + K_z \Pi.$$

#### C.4. Kalman Filter

The states $z[k]$ and $w[k]$ of the underlying system are needed to compute the control action in Eq. (S16). We compute minimum-variance estimates for these states from observed intensity measurements by implementing a Kalman filter on the combined system dynamics in ( Eq. (S14), Eq. (S15)). The dynamics of both $z[k]$ and $w[k]$ combine into an augmented state $\xi := \left[z^\top \; w^\top\right]^\top \in \mathbb{R}^8$, with combined dynamics:

$$\begin{aligned} \xi[k+1] &:= \begin{bmatrix} z[k+1] \\ w[k+1] \end{bmatrix} = \overbrace{\begin{bmatrix} A & B_{\text{lqr}} \\ 0 & S \end{bmatrix}}^{A_f} \xi[k] + \overbrace{\begin{bmatrix} B \\ 0 \end{bmatrix}}^{B_f} u_{\text{lin}}[k] + \nu_f[k], \\ y_a[k] &:= \begin{bmatrix} y_{\text{lin}}[k] \\ r[k] \end{bmatrix} = \underbrace{\begin{bmatrix} C_z & C_{\text{lqr}} \\ 0 & C_r \end{bmatrix}}_{C_f} \xi[k] + \mu_f[k], \end{aligned}$$

where the augmented measurement vector $y_a[k] \in \mathbb{R}^2$ includes both the measurement deviation $y_{\text{lin}}[k]$ and the reference trajectory $r[k]$. The augmented noise processes $\nu_f[k] \sim \mathcal{N}(0, Q_f)$ and $\mu_f[k] \sim \mathcal{N}(0, R_f)$ are assumed to be Gaussian in the Kalman filter design, with covariance matrices $Q_f$ and $R_f$. This assumption works well in practice, even though they do not represent the exact noise processes in the underlying system.

The Kalman estimator predicts the state at the next time-step ($\hat{\xi}[k+1|k]$) after initialisation of the state $\hat{\xi}[0|0]$ and the covariance $P[0|0]$. This prediction update is a direct evaluation of the system dynamics under an input $u_{\text{lin}}[k]$,

$$\begin{aligned} \hat{\xi}[k+1|k] &= A_f \hat{\xi}[k|k] + B_f u_{\text{lin}}[k], \\ P[k+1|k] &= A_f P[k|k] A_f^\top + Q_f. \end{aligned} \tag{S19}$$

Upon acquiring the next measurement $y[k]$ and reference target $r[k]$, the filter computes the optimal Kalman gain $K_k \in \mathbb{R}^{8\times 2}$ to correct the prediction for the estimate $\hat{\xi}[k|k]$:

$$\begin{aligned} K_k &= P[k|k-1] C_f^\top \left(C_f P[k|k-1] C_f^\top + R_f\right)^{-1}, \\ \hat{\xi}[k|k] &= \hat{\xi}[k|k-1] + K_k \left(y_a[k] - C_f \hat{\xi}[k|k-1]\right), \\ P[k|k] &= (I - K_k C_f) P[k|k-1]. \end{aligned} \tag{S20}$$

Alternating the prediction Eq. (S19) and measurement update Eq. (S20) yields an optimal, minimum-variance estimate for $\hat{\xi}[k]$, with covariance $P[k]$.

Typically, the matrices $Q_f$ and $R_f$ are constructed by hand through expert and sensor knowledge, which are unavailable for biological applications. Therefore, we compare strategies of heuristic tuning and optimised tuning for the state covariance, while computing a numerical estimate for $R_f$ from residual data $e[k] = y[k] - \hat{y}[k]$. The variance of these residuals led to $R_f = \operatorname{diag}(0.02, 10^{-7})$, with the predictions $\hat{y}[k]$ computed by simulation of the individual models. The second entry was chosen manually to reflect low uncertainty in the reference trajectory. In practice, the Kalman filter covariance matrices are frequently considered as tuning parameters rather than true covariance matrices. In the context of filtering for nonlinear systems, optimised covariance matrices even allow linear filters to achieve tracking performance similar to neural-network-based methods [14].

We first consider the heuristic tuning of the process noise covariance $Q_f$, which balances the confidence of the estimator in the model and the incoming measurements. High model confidence was enforced by setting the corresponding diagonal elements to $Q_{f,ii} = 10^{-4}$ for $i = 1 \dots 3$. Conversely, the estimator should adapt rapidly to prediction errors through $w[k]$, but not overcorrect to measurement noise. This rapid adaptation was achieved by setting $Q_{f,44} = 0.01$ and $Q_{f,55} = 0.001$. Finally, even faster adaptation is required for the reference-tracking states, yielding $Q_{f,ii} = 10^{-1}$ for $i = 6 \dots 8$.

To obtain the optimised tuning matrices, we minimised both the prediction residual between the predicted $\hat{x}_p[k|k-1]$ and simulated states $x_p[k]$, and the residual between predicted measurements $\hat{y}_p[k|k-1]$ and experiment measurements $y_p[k]$. We chose this strategy because predictions are more directly affected by the covariance $Q_f$ (Eq. (S19)). The index $p$ indicates the traces used during optimisation of the filter parameters. To reduce optimisation time, we took a random sample of $N_p = 10$ traces from the system identification dataset. The complete optimisation problem was formulated and solved using Matlab's `patternsearch` solver [15],

$$\begin{aligned}
&\min_{Q} \sum_{p=0}^{N_p-1} \sum_{k=1}^{N-1} \|T\hat{z}_p[k|k-1] + x^* - x_p[k]\|_2^2 + \lambda \|\hat{y}_p[k|k-1] - y_p[k]\|_2^2 \\
\text{s.t.} \quad & x_p[j+1] \quad = F(x_p[j], u_p[j]), \\
& x_p[0] \quad = x_p(t_0) \qquad\qquad k = 0, 1, \dots, N-1 \\
& \hat{\xi}_p[k+1|k] \quad (\text{Eq. (S19)}, \text{Eq. (S20)}) \qquad p = 0, 1, \dots, N_p - 1 \\
& \hat{y}_p[k+1|k] \quad = \begin{bmatrix} C & C_{\text{lqr}} \end{bmatrix} \hat{\xi}_p[k+1|k] + y^* \\
& Q \quad \succeq 0
\end{aligned} \tag{S21}$$

Here, the estimate $\hat{z}[k|k-1]$ indicates the estimate for Eq. (S15), derived from the augmented state vector $\hat{\xi}[k|k-1]$. The transition function $F(x_p[j], u_p[j])$ indicates the discretised dynamics of the aggregate model in Eq. (S10). The initial condition $x_p(t_0)$ used for this simulation was the initial condition found during system identification for that trace. Finally, we selected a balancing parameter $\lambda = 100$ to prioritise minimisation of the output-prediction residual.

No direct experimental measurements of the internal active and inactive cytosolic or nucleic concentrations are available. Therefore, a simulation of the individual models was used to compare Kalman filter performance (Fig. S3). We compare the state and output estimates produced by the heuristically tuned Kalman filter with those produced from the optimised Kalman filter in Figure S3. In addition, the absolute error $|\hat{x} - x|$ and $|\hat{y} - y|$ are presented. We used the state transformation from section C.1 to recover the concentration states in Eq. (S1),

$$\hat{x}[k] = T \begin{bmatrix} 0 \\ z[k|k] \end{bmatrix} + x^*.$$

The results in Figure S3 indicate that estimates obtained from either method have a similar absolute error. Therefore, the heuristically tuned Kalman filter was preferred for brevity and simplicity in the main article.

To summarise, each time we take a measurement, we estimate $z[k]$ and $w[k]$ using the Kalman filter equations Eq. (S19) and Eq. (S20). Then, the control action is computed using the feedback law Eq. (S16), with the gains $K_w$ and $K_z$ computed offline.

### D. Model Predictive Control

MPC evaluates the optimisation problem numerically, directly accommodating nonlinear and continuous-time dynamics. We therefore formulate the control problem directly with the continuous-time dynamics in Eq. (S1), leveraging the numerical integration schemes in the `do-mpc` package to handle discretisation. The aggregate physics-based model was again used to model this controller.

Notation in this section will reflect the similarities between LQR and MPC to maintain clarity. We explicitly define the changes in (virtual) model equations, while keeping the notation similar when variables play the same role. These variables were previously indexed with 'lqr' and will be indexed with 'mpc' in this section. The EKF operates on the concentration state ($x(t) \in \mathbb{R}^4, u(t) \in \mathbb{R}_+$) instead of the linearised and truncated state ($z[k], u_{\text{lin}}[k]$).

Similar to LQR, we counteract unmodeled cell-to-cell variations and external disturbances by augmenting the controller with a virtual system. This virtual disturbance $w_{\text{mpc}} \in \mathbb{R}^2$ was modeled,

$$\frac{\mathrm{d}}{\mathrm{d}t} w_{\text{mpc}}(t) := \frac{\mathrm{d}}{\mathrm{d}t} \begin{bmatrix} d_x(t) \\ d_y(t) \end{bmatrix} = \underbrace{\begin{bmatrix} -0.01 & 0 \\ 0 & -0.01 \end{bmatrix}}_{S_{\text{mpc}}} \begin{bmatrix} d_x(t) \\ d_y(t) \end{bmatrix}$$

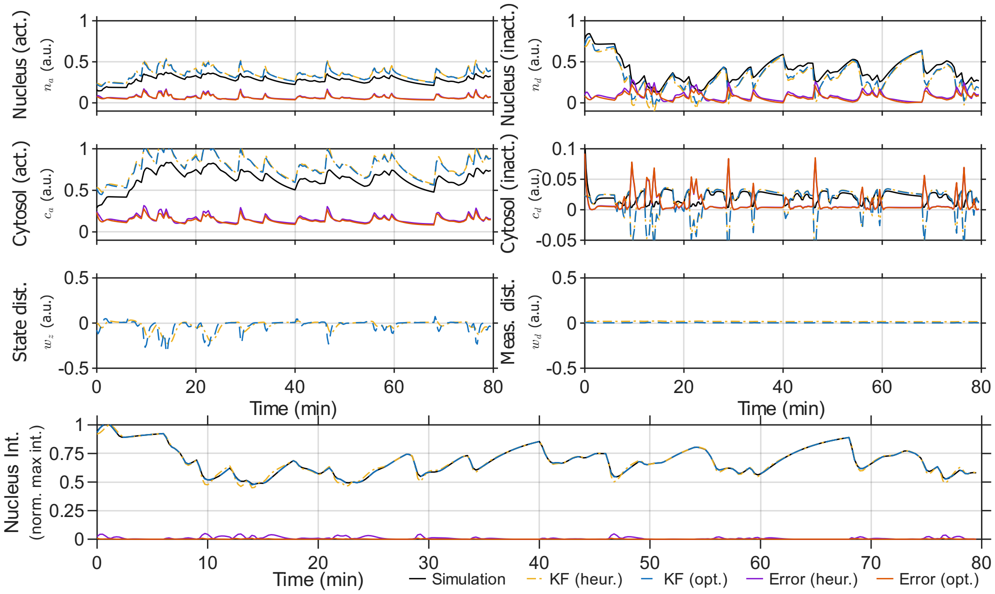


**Fig. S3.** Internal states, output and absolute error of individual model simulation compared to the state and output estimated by the Kalman filter. Both for a filter tuned by hand using heuristic rules (heur.), and tuned using an optimisation procedure (opt.).

where the diagonal entries of $S_{\mathrm{mpc}} \in \mathbb{R}^2$ again represent near-constant disturbances, with a small decay to aid controller stability. We introduced the notation for $d_x$ and $d_y$ to indicate how these disturbances affect the augmented state-space model. This augmented model combines the continuous-time dynamics of the system with the virtual model,

$$\frac{\mathrm{d}}{\mathrm{d}t}x(t) = f_n(x(t), u(t)) + B_{\mathrm{mpc}} w_{\mathrm{mpc}}(t)$$
$$y(t) = Cx(t) + C_{\mathrm{mpc}} w_{\mathrm{mpc}}(t),$$

where $x(t) \in \mathbb{R}^4$ reflects the nuclear and cytosolic concentrations, $u(t) \in \mathbb{R}$ the activation irradiance and $y(t)$ the normalised intensity measurements. The input disturbance $B_{\mathrm{mpc}}$ was designed to affect the photo-activation rate at the same rate as the optogenetic input ($k_n$ in Eq. (S1)). In the main text we demonstrated a variation in the dark-state activation, motivating this choice. The output disturbance is a static measurement bias,

$$B_{\mathrm{mpc}} = \begin{bmatrix} k_n & 0 \\ -k_n & 0 \\ 0 & 0 \\ 0 & 0 \end{bmatrix}, \qquad C_{\mathrm{mpc}} = \begin{bmatrix} 0 & 1 \end{bmatrix}.$$

We now formulate the mixed discrete-time and continuous-time optimisation problem used for MPC. Given the sampling interval of $\tau = 15$ s and a prediction horizon $N_h = 10$, the complete optimisation problem that is solved at time-step $k$ becomes

$$\begin{aligned}
\min_{\substack{u(j\tau),\ j=k\dots k+N_h}} \quad & \sum_{j=k}^{k+N_h} Q_{\mathrm{mpc}}(y(j\tau)-r[j])^2 + R_{\mathrm{mpc}}u^2(j\tau) + W(u(j\tau)-u((j+1)\tau))^2 \\
\text{s.t.} \quad & \dot{x}(t) = f_n(x(t),u(t)) + B_{\mathrm{mpc}}w_{\mathrm{mpc}}(t), \\
& \dot{w}_{\mathrm{mpc}}(t) = S_{\mathrm{mpc}}w_{\mathrm{mpc}}(t), \\
& y(t) = Cx(t) + C_{\mathrm{mpc}}w_{\mathrm{mpc}}(t), \\
& u(t) = u(t_j) \qquad t_j \in [j\tau,(j+1)\tau), \qquad j = k,\dots,k+N_h-1, \\
& x(k\tau) = x_0, \\
& w_{\mathrm{mpc}}(k\tau) = w_0, \\
& u(t) \in [0, 5.8].
\end{aligned} \tag{S22}$$

The tracking cost $Q_{\mathrm{mpc}} \in \mathbb{R}_+$, the control effort penalty $R_{\mathrm{mpc}} \in \mathbb{R}_+$ and the input rate penalty $W \in \mathbb{R}_+$ balance the trade-off between reference tracking and smooth actuator performance. The main text outlines the tuning procedure used. The MPC problem is initialised with $w_0$ and $x_0$, which are state estimates obtained from an extended Kalman filter. Finally, the optimisation problem was implemented and executed with the `do-mpc` Python package, which uses Casadi to formulate symbolic optimisation equations and the IPOPT interior-point solver to solve the optimisation problem [16–18].

##### D.1. Extended Kalman filter

The MPC formulation in Eq. (S22) relies on estimates for the initial states $x_0$ and $w_0$ at timestep $k$. Similar to LQR, we estimate these initial states using an extended Kalman filter (EKF) applied to a discretisation of the LEXY dynamics [19]. The EKF is a nonlinear analogue of the classical Kalman filter, replacing the update equations with their nonlinear counterparts. While global stability of the EKF is not guaranteed, the weak nonlinearity of the transport equation is expected to lead to convergence in our application.

State propagation follows a minor adaptation of the nonlinear dynamics due to solution simplicity. The disturbance effect was modeled as a separately integrated contribution,

$$\hat{\xi}_{\mathrm{ekf}}[k+1|k] := \begin{bmatrix} \hat{x}[k+1|k] \\ \hat{w}_{\mathrm{mpc}}[k+1|k] \end{bmatrix} = \begin{bmatrix} F(\hat{x}[k|k],u[k]) + \tau B_{\mathrm{mpc}}\hat{w}_{\mathrm{mpc}}[k|k] \\ \exp\left(S_{\mathrm{mpc}}\tau\right)\hat{w}_{\mathrm{mpc}}[k|k] \end{bmatrix}, \tag{S23}$$

where the function $F$ is the discretisation computed in S10. The corresponding covariance propagation is computed from the Jacobian of the dynamics $F_k \in \mathbb{R}^{6\times 6}$, linearised at the current operating point,

$$\begin{aligned}
F_k &= \begin{bmatrix} \frac{\partial F}{\partial x}\big|_{\hat{x}[k|k],u[k]} & B_{\mathrm{mpc}} \\ 0 & S_{\mathrm{mpc}} \end{bmatrix}, \\
P_{\mathrm{ekf}}[k+1|k] &= F_k P_{\mathrm{ekf}}[k|k] F_k^\top + Q_{\mathrm{ekf}}.
\end{aligned} \tag{S24}$$

The measurement update equations are equivalent to that of the original Kalman filter (Eq. (S20)), but with the output matrix $C_f = [C \;\; C_{mpc}]$. The interpretation of the state covariance matrix $Q_{\mathrm{ekf}} \in \mathbb{R}_+^{6\times 6}$ and $R_{\mathrm{ekf}} \in \mathbb{R}_{>0}$ remain equivalent to that of the linear Kalman filter.

Similar to the linear Kalman filter, no exact results for the covariances $Q_{\mathrm{ekf}}$ and $R_{\mathrm{ekf}}$ can be obtained. Instead, we again compare reconstruction accuracy between heuristic and optimised tuning for the EKF. In both cases, we use the estimate for $R_{\mathrm{ekf}} = 0.02$ obtained from measurement data. For the state covariance $Q_{\mathrm{ekf}}$ we used the same covariance estimates as with the linear Kalman filter, setting $Q_{\mathrm{ekf},ii} = 10^{-4}$ for $i = 1\dots 4$, $Q_{\mathrm{ekf},55} = 0.01$ and $Q_{\mathrm{ekf},66} = 0.001$.

The optimisation problem used for optimal tuning is equivalent to that of the linear Kalman filter with the nonlinear update equations,

$$
\begin{aligned}
&\min_{Q} \sum_{p=0}^{N_p-1} \sum_{k=1}^{N-1} \|\hat{x}_p[k|k-1] - x_p[k]\|_2^2 + \lambda \|\hat{y}_p[k|k-1] - y_p[k]\|_2^2 \\
\text{s.t.}\quad & x_p[j+1] &&= F(x_p[j], u_p[j]), \\
& y_p[j] &&= h_n(x_p[j]), \\
& x_p[0] &&= x_p(t_0) && k = 0, 1, \ldots, N-1 \\
& \hat{\xi}_p[k+1|k] && \qquad (\text{Eq. (S23)},\ \text{Eq. (S24)}) && p = 0, 1, \ldots, N_p - 1 \\
& \hat{y}_p[k+1|k] &&= \begin{bmatrix} C & C_{\text{mpc}} \end{bmatrix} \hat{\xi}_p[k+1|k] \\
& Q_{\text{ekf}} &&\succeq 0.
\end{aligned}
\tag{S25}
$$

Again, the index $p$ is used to refer to the different traces used during model identification, sampling $N_p = 10$ random traces. Recall that $\hat{x}[k|k-1]$ can be extracted from the combined state estimate $\hat{\xi}[k|k-1]$. We solve this optimisation problem using the same approach as for the linear Kalman filter, with the same initial condition $x_p(t_0)$ and balancing parameter $\lambda = 100$ (Section C.4).

The comparison in Figure S4 compares the state prediction under heuristic and optimised tuning. The difference between both methods is small, with the two approaches leading to only slight differences in protein concentrations. Therefore we used the heuristically tuned EKF in the main paper to demonstrate indicative performance.

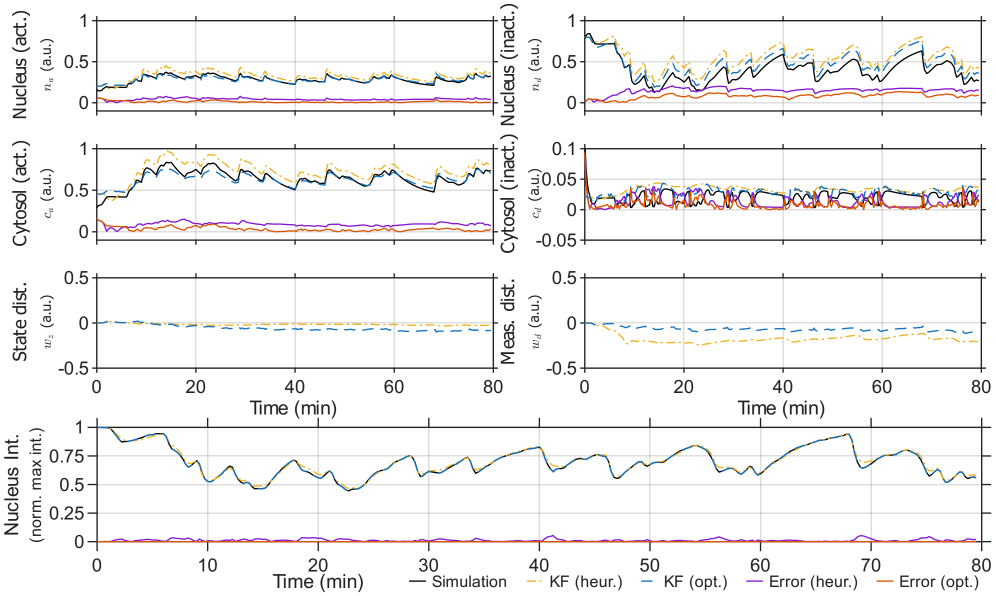


**Fig. S4.** Internal states, output and absolute error of individual model simulation compared to the state and output estimated by the EKF. Both for a filter tuned by hand using heuristic rules (heur.), and tuned using an optimisation procedure (opt.).
